# The N-terminal subunit of vitellogenin in planthopper eggs and saliva acts as a reliable elicitor that induces defenses in rice

**DOI:** 10.1101/2022.05.24.493233

**Authors:** Jiamei Zeng, Wenfeng Ye, Wenhui Hu, Xiaochen Jin, Peng Kuai, Wenhan Xiao, Yukun Jian, Ted C. J. Turlings, Yonggen Lou

**Affiliations:** State Key Laboratory of Rice Biology & Ministry of Agriculture Key Lab of Molecular Biology of Crop Pathogens and Insects, Institute of Insect Sciences, Zhejiang University, Hangzhou 310058, China; Laboratory of Fundamental and Applied Research in Chemical Ecology, Institute of Biology, University of Neuchâtel, Neuchâtel 2000, Switzerland

## Abstract

Vitellogenins are essential for the development and fecundity of insects, but these proteins may also betray them, as we show here. We found that the small N-terminal subunit of vitellogenins of the planthopper *Nilaparvata lugens* (NlVgN) triggers strong defense responses in rice plants when it enters the plant during feeding or oviposition by the insect. The defenses induced by NlVgN in plants not only decreased the hatching rate of *N. lugens* eggs, but also induced volatile emissions in rice plants, which rendered them attractive to a common egg parasitoid. VgN of other planthoppers were found to trigger the same defense responses in rice. We further show that VgN deposited during planthopper feeding compared to during oviposition induces a somewhat different response, probably targeting the appropriate developmental stage of the insect. The key importance of VgN for planthopper performance precludes possible evolutionary adaptions to prevent detection by rice plants.

## Introduction

When attacked by herbivores, plants perceive elicitors derived from herbivores and then activate early signaling events, such as the increase in concentrations of cytosolic calcium ion (Ca^2+^), the activation of mitogen-activated protein kinases (MAPKs) and the production of reactive oxygen species (ROS) (Wu and Baldwin, 2010). The early signaling events lead to the activation of signaling pathways mediated by defense-related phytohormones, which mainly consist of jasmonic acid (JA), salicylic acid (SA) and ethylene (ET). The activated signaling pathways mediate the production of defensive compounds and thus enhance the resistance of plants to herbivores (Schuman and Baldwin, 2016).

Several elicitors, such as fatty acid-amino acid conjugates (FACs), inceptins, caeliferins, bruchins, benzyl cyanide, and indole, have been identified in oral secretions, oviposition fluids and feces of herbivores (Hilker and Fatouros, 2015; Ray et al., 2016; Chen and Mao, 2020). These elicitors are mostly herbivore species-specific and can induce targeted defense responses in plants (Arimura, 2021). So far, elicitors are mainly known for chewing herbivores (Chen and Mao, 2020), but a few have also been identified for piercing-sucking herbivores, such as phosphatidylcholines isolated from female white-backed planthoppers (WBPH, *Sogatella furcifera*), a bacterial chaperonin GroEL from the saliva of potato aphid (*Macrosiphum euphorbiae*), and a mucin-like protein from the saliva of the brown planthopper (BPH, *Nilaparvata lugens* (Stål)) (Chaudhary et al., 2014; Yang et al., 2014; Shangguan et al., 2018). These cases almost exclusively involve elicitors in the insects’ oral secretions, but it is known that egg deposition may also activate plant defense responses (Hilker and Fatouros, 2015). Certain insect-derived compounds have been implicated in such oviposition-related responses (Reymond, 2013; Gouhier-Darimont et al., 2013, 2019; Hilker and Fatouros, 2015; Bertea et al., 2020), but to date, only phosphatidylcholines have been identified as specific egg-derived elicitors (Stahl et al., 2020).

Vitellogenins (Vgs) are the major yolk protein precursors that are vital for the egg development in most oviparous vertebrate and invertebrate animals (Tufail and Takeda, 2008). Insect Vgs are mostly synthesized by the fat body in a sex-, stage- and tissue-specific manner (Raikhel and Dhadialla, 1992). After synthesis in the fat body, Vgs are typically cleaved into two subunits, a small N-terminal subunit (< 65 kDa; VgN) and a large C-terminal subunit (> 150 kDa) at a consensus cleavage site, R/K-X-R/K-R, by subtilisin-like endoproteases (Tufail and Takeda, 2008). They are then structurally modified, such as proteolytic cleavage, glycosylation, phosphorylation and lipidation, and subsequently secreted into the hemolymph and taken up by oocytes via receptor-mediated endocytosis (Tufail et al., 2014). Insect Vgs were previously considered as female-adult-specific proteins and only produced in fat body cells. However, recent studies have revealed that Vgs are also found in sexually immature individuals and male adults (Piulachs et al., 2003; Huo et al., 2018). Moreover, *Vg* genes have been found abundantly expressed in hemocytes and ovaries, in addition to fat bodies (Chen et al., 2012; Huo et al., 2018). In some species, albeit at low levels, they are also expressed in salivary glands, midguts and non-neuronal glial cells (Münch et al., 2015; Shen et al., 2019). Consistent with their distributions in insect bodies, multiple non-nutritional functions have been attributed to Vgs, in addition to their nutritional functions. The Vg from honey bee *Apis mellifera* affects food-related behaviors and some survival traits such as immunity, oxidative stress resilience and lifespan (Amdam et al., 2012), as well as the transport of immune elicitors from mother to offspring (Salmela et al., 2015). The Vg of mosquito *Anopheles gambiae* is capable of interfering with the anti-*Plasmodium* response (Rono et al., 2010). Insect Vgs are also involved in the vertical transmission of plant viruses by binding to viral proteins (Wei et al., 2017; Huo et al., 2018). The key role that Vgs play in important physiological processes in insects, as well as their specific chemical features, make them susceptible to recognition by other organisms.

The brown planthopper (BPH) (Hemiptera: Delphacidae), a monophagous piercing-sucking herbivore, is one of the most important insect pests of rice (*Oryza sativa* L.) in Asia (Dyck and Thomas, 1979). It damages plants by feeding on phloem sap (causing minor tissue damages via its stylet), laying egg clusters in tissues (causing more tissue damages via its ovipositor), but harm is also caused by viruses transmitted by BPH (Sōgawa, 1982; Hattori and Sōgawa, 2002). Newly-emerged BPH female adults do not lay eggs until they pass their pre-oviposition periods, which generally takes about three days at 25-28 ℃ (Mochida and Okada, 1979). The mature Vg of BPH, NlVg, contains two vitellogenin-N domains at N-terminus, a middle-region domain of unknown function (DUF1943) and a von Willebrand factor type D (vWD) domain at C-terminus (Tufail et al., 2010). The predicted molecular weight of NlVg is 227.94 kDa, and the mature protein is typically cleaved into a small N-terminal subunit (48.33 kDa, NlVgN) and a large C-terminal subunit (179.24 kDa) at the RSRR sequence motif of the N-terminus (Cheng and Hou, 2005; Tufail et al., 2010). In addition to high transcript levels in the fat body of BPH female adults (Tufail et al., 2010), *NlVg* was observed to be abundantly expressed in salivary glands of adult females (Noda et al., 2008; Ji et al., 2013). Moreover, *NlVg* is also expressed in head, midgut, epidermis and thorax of female adults, midgut and testes of male adults, salivary glands of 3^rd^ to 5^th^ instar nymphs, and whole body of eggs, 1^st^ to 5^th^ instar nymphs, female adults and male adults (Wang et al., 2015; Shen et al., 2019). NlVg plays an important role in BPH growth, development, and fecundity; knockdown of *NlVg* causes abnormal increases in body size and mass of BPH female adults, but reduces the number of eggs laid by female adults (Shen et al., 2019). In BPH, there are also two NlVg-like genes, NlVg-like1 and NlVg-like2, both of which are not clustered with the conventional insect Vgs, including NlVg (Shen et al., 2019). *NlVg* and *NlVg*-like2 exhibit similar expression patterns in BPH developmental stages and tissues, whereas *NlVg-like1* shows different patterns. Although each of the three genes influences BPH egg development and fecundity, their specific functions appear to differ (Shen et al., 2019).

NlVgN exists not only in BPH hemolymph but also in gelling saliva of 5^th^ instar nymphs, as well as in their eggs and oviposition fluids (Tufail et al., 2010; Xie, 2012; Huang et al., 2016). During feeding the saliva enters rice tissues and coagulates to form a salivary sheath around the stylets, whereas during oviposition fluids are deposited to glue the eggs to the damaged plant tissue. This implies that significant amounts of NlVgN will enter rice tissues when BPH infests plants. It has been reported that defense responses in rice induced by BPH gravid female infestation are distinctly different from those induced by nymphal infestation. For example, infestation by gravid BPH females enhances levels of JA and JA-Ile but decreases ethylene levels in rice, whereas infestation by BPH nymphs does not change the levels of these phytohormones (Li, 2015; Ji et al., 2017; Ye et al., 2017). These distinct responses prompted us to hypothesize that NlVgN in BPH saliva and eggs plays a role in regulating the interaction of adult BPH with rice.

In this study, we tested our hypothesis by exploring the role of *NlVg* (GenBank: AB353856) in BPH-induced defense responses in rice. Combining molecular tools, chemical analyses and bioassays, we revealed that NlVgN can indeed enter rice plants via the BPH saliva or from the surface of BPH eggs. We show that, together with the damage caused by BPH feeding or oviposition, the small N-terminal subunit of NlVg allows rice plants to specifically recognize planthopper attacks. As this protein is essential for planthopper survival, it is seemingly impossible for the planthoppers to avoid this recognition.

## Results

### NlVgN enters rice tissues during BPH feeding and oviposition

BPH causes damage to rice plants by sucking phloem sap or laying its eggs in rice tissues (Figure 1A-D). As is the case for other insects, the eggs of BPH contain high levels of vitellogenins (Cheng and Hou, 2005; Tufail et al., 2010), but it has been found that NIVgs, especially NlVgN, are also present in BPH gelling saliva (Huang et al., 2016; Rao et al., 2019). We therefore examined whether NlVgN enters rice tissues during BPH feeding as well as oviposition.

**Figure 1.**
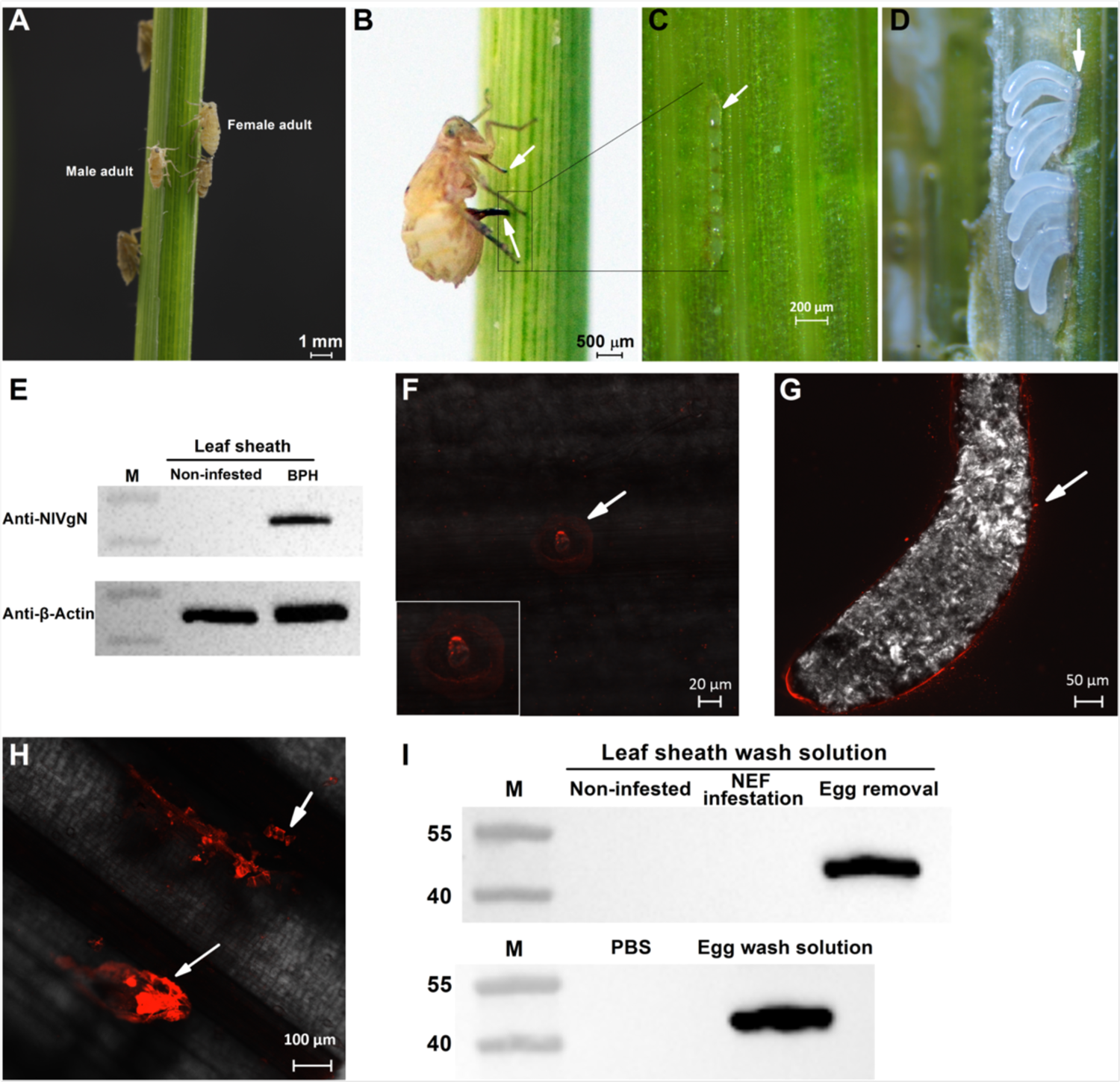
NlVgN enters rice tissues during BPH feeding and egg deposition. (**A**) A rice plant exposed to BPH infestation, showing male and female adults. (**B**) A gravid BPH female feeding (see upper arrow) and ovipositing (see lower arrow) on a rice plant. (**C**) BPH eggs in the rice leaf sheath with the arrow pointing out the egg caps. (**D**) BPH eggs are visible after the leaf sheath was removed. The upper approximately transparent part of the egg, indicated by the arrow, is the egg cap. (**E**) Western blot showing that NlVgN is present in rice plants that were infested for 24 h by 50 newly-emerged BPH female adults, but not in non-infested plants. (**F**-**H**) Immunofluorescence microscopy reveals the presence of NlVgN (indicated by the arrow) at BPH feeding sites (**F**), on the surface of the egg (**G**) and in egg caps (**H**). (**I**) Western blot showing that NlVgN is present in the PBS buffer in which pieces of leaf sheaths that were infested by BPH gravid females but BPH eggs were removed (Egg removal) and intact BPH eggs (Egg wash solution) were immersed for 8-10 min and 2-3 min, respectively, but not in the buffer in which pieces of leaf sheaths that were kept non-infested or infested by BPH Newly Emerged Females (NEF infested) were immersed for 8-10 min or in the buffer alone. M, molecular weight markers (kDa).

Western blot analysis with anti-NlVgN showed a band of about 50 kDa in total proteins from rice leaf sheaths that had been infested by newly-emerged BPH females (feeding only) for 24 h, whereas no NlVgN band was detected in proteins from non-infested plants (Figure 1E). Consistent with this finding we also found, with an immunofluorescence assay (IFA) using anti-NlVgN, that NlVgN is present around BPH feeding sites (Figure 1F), implying that NlVg indeed enters rice tissues when BPH feeds on plants.

NlVgN also occurs on the surface of BPH eggs (Figure 1G) and specifically in the egg cap (Figure 1H), which is the upper somewhat transparent part of the egg that adheres to oviposition-damaged rice tissues (Figure 1C and D). Additionally, a NlVgN band was detected in the PBS buffer solution used to extract pieces of rice leaf sheaths on which BPH females had oviposited but from which eggs had been carefully removed. The band was also found in similar extracts from intact BPH eggs, whereas no band was detected in PBS solutions used to extract pieces of rice leaf sheaths infested with newly-emerged BPH females (no eggs) or uninfested leaf sheaths (Figure 1I). These results confirm that NlVgN on the surface of BPH eggs come in contact with damaged rice tissues during oviposition. Taken together, these results suggest that NlVgN reaches rice plants during BPH feeding and oviposition.

### NlVgN induces the production of cytosolic Ca^2+^ and H_2_O_2_ in rice

It has been reported that BPH nymphs, which only feed, as well as gravid females, which feed and lay eggs, induce increases in the concentration of cytosolic Ca^2+^ and H_2_O_2_ in rice plants. Both of these responses affect signaling pathways that play an important role in rice defenses (Zhou et al., 2009; Ye et al., 2017). To explore the role of NlVgN plays in activating these two pathways, we silenced the *NlVg* gene in BPH using RNA interference (RNAi) as described previously (Liu et al., 2010) and then investigated the effect of *NlVg* silencing on the levels of cytosolic Ca^2+^ and H_2_O_2_ in rice. Injecting fifth instar nymphs with double-stranded RNA (dsRNA) of the N-terminal sequence of *NlVg* (*dsNlVg*) reduced the transcript level of *NlVg* in BPH female adults by 97.90, 90.64 and 73.06% at 1, 3, and 5 d after emergence (2-6 d post injection), respectively, compared to those in BPH injected with double-stranded RNA of green fluorescent protein (*dsGFP*) (Figure 2—figure supplement 1A). The protein level of NlVgN also decreased drastically in BPH injected with *dsNlVg* (dsNlVg-BPH) (Figure 2—figure supplement 1B). This dsRNA injection did not co-silence *NlVg-like1* and *NlVg-like 2*, showing that the RNAi is specific (Figure 2—figure supplement 1C and 1D). Compared to the levels in plants infested by newly-emerged dsGFP-BPHs adult females (feeding only), the level of NlVgN in plants infested by newly-emerged dsNlVg-BPHs adult female was also considerably lower (Figure 2A). Cytosolic Ca^2+^ and H_2_O_2_ analysis revealed that feeding by dsNlVg-BPHs, compared with feeding by dsGFP-BPHs and C-BPHs, induced a weaker fluorescence intensity around feeding sites at 3 h (Figure 2B and C) and lower levels of H_2_O_2_ at 3 and 8 h after BPH infestation (Figure 2D). Moreover, application of recombination protein NlVgN increased the H_2_O_2_ level in rice plants 15-30 min after treatment (Figure 2E and F), whereas expressing *NlVgN* in rice (Figure 2—figure supplement 2) enhanced constitutive and BPH-induced (8 h after infestation) levels of H_2_O_2_ in plants (Figure 2G). These findings demonstrate that NlVgN, either secreted by BPH salivary glands or on the surface of BPH eggs, contributes to BPH-induced increases of cytosolic Ca^2+^ and production of H_2_O_2_ in rice.

**Figure 2.**
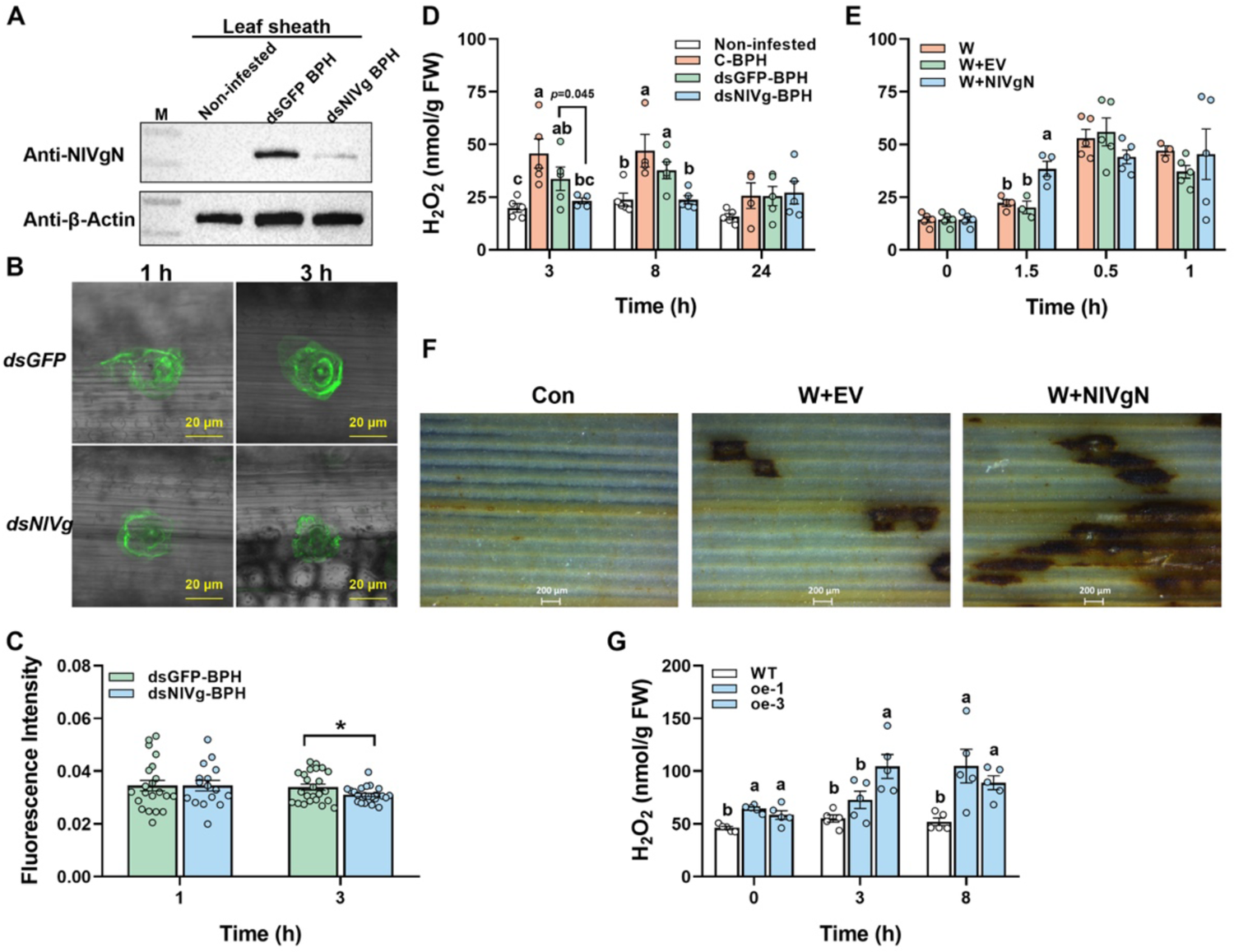
NlVgN enhances BPH feeding-induced concentrations of cytosolic Ca^2+^ and H_2_O_2_ in rice. (**A**) Detection of protein fraction NlVgN in rice infested for 24 h by 50 newly-emerged BPH female adults (12 to 24 h after emergence) that had been injected with dsRNA of *GFP* (*dsGFP*) or *NlVg* (*dsNlVg*) at fifth-instar nymph stage, or kept non-infested. M indicates molecular weight markers. (**B** and **C**) Confocal microscopic images showing green fluorescence of Fluo-3 AM binding with intracellular Ca^2+^ at BPH feeding sites (indicated by the arrow) of rice leaf sheaths (**B**) and mean fluorescence intensity (+SE, n = 16-24) at feeding sites (**C**) that were infested for 1 and 3 h by newly-emerged female adults that had been injected with *dsGFP* or *dsNlVg* at fifth-instar nymph stage. Asterisk indicates significant difference between treatments (\**P* < 0.05, Student’s *t*-test). (**D**) Mean levels (+SE, n = 5) of H_2_O_2_ in leaf sheaths of TN1 plants that were kept non-infested (Non-infested) or infested for 3, 8 and 24 h by newly-emerged BPH female adults that had been injected with *dsGFP*, *dsNlVg* or kept non-injected (C-BPH) at fifth-instar nymph stage. (**E**) Mean levels (+SE, n = 5) of H_2_O_2_ in leaf sheaths of TN1 plants that were kept unmanipulated (0 h) or treated for 0.5, 1 and 3 h with wounding plus the purified recombinant protein NlVgN (W+NlVgN), the purified products of the empty vector (EV) (W+EV) or nothing. (**F**) In situ detected H_2_O_2_ accumulation in rice leaves by 3,3’-diaminobenzidine (DAB) staining. Plant leaves were kept unmanipulated (Con) or treated for 15 min with W+EV or W+NlVgN. (**G**) Mean levels (+SE, n = 5) of H_2_O_2_ in XS11 plants expressing *NlVgN* (line oe-1 and oe-3) and wild-type (WT) plants that were kept non-infested or infested with 10 gravid BPH female adults for 3, 8 and 24 h. FW, fresh weight. Letters indicate significant differences among different treatments (*P* < 0.05, Tukey’s HSD post-hoc test).

### Silencing *NlVg* does not affect the production of JA and JA-Ile in rice fed on by BPH but exogenous NlVgN or expressing *NlVgN* in rice does

JA- and ethylene-mediated signaling pathways play a central role in regulating the resistance of rice to BPH (Zhou et al., 2009; Lu et al., 2014; Xu et al., 2021). Moreover, infestation by gravid BPH females (feeding + oviposition) induces the production of JA and JA-Ile and inhibits the production of ethylene in rice, whereas infestation by nymphs does not (Li, 2015; Ji et al., 2017; Ye et al., 2017; Ye et al., 2020; Xu et al., 2021). Hence, we wondered if NlVgN plays a role in modulating the biosynthesis of these phytohormones in rice. Consistent with previous results (Ji et al., 2017), BPH feeding did not induce the production of JA and JA-Ile, nor did the knockdown of *NlVg* affect JA and JA-Ile levels after BPH feeding. In each case hormone production was very low (Figure 3—figure supplement 1A and 1B). Although wounding plus EV resulted in higher JA and JA-Ile levels than in unmanipulated plants, the treatment with wounding plus purified recombination protein NlVgN induced still higher levels of JA and JA-Ile than in plants with wounding plus EV (Figure 3A and B). Consistent with these results, wounding plus either the homogenized fresh BPH egg solution in a phosphate buffer (pH 7.4), the homogenized BPH eggshell solution in the buffer, or the homogenized dsGFP-BPH ovary solution in the buffer, all significantly induced the biosynthesis of JA and JA-Ile in plants compared to wounding plus the buffer alone (Figure 3C-F), whereas wounding plus the homogenized dsNlVg-BPH ovary solution in the buffer (low levels of NlVgN) exhibited impaired induction of JA and JA-Ile (Figure 3E and F). Additionally, plants expressing *NlVgN* showed high constitutive and BPH-induced (infestation for 24 h) levels of JA and JA-Ile (Figure 3G and H). Treatment with wounding plus the recombination protein NlVgN did not induce the biosynthesis of ethylene (Figure 3—figure supplement 1C). Taken together, these findings indicate that NlVgN-induced production of JA and JA-Ile in rice is dependent on damage level or type of the tissue that NlVgN comes in contact with and/or on the effectors and other elicitors derived from BPH feeding or oviposition. These data further show that NlVgN does not affect the biosynthesis of ethylene in rice.

**Figure 3.**
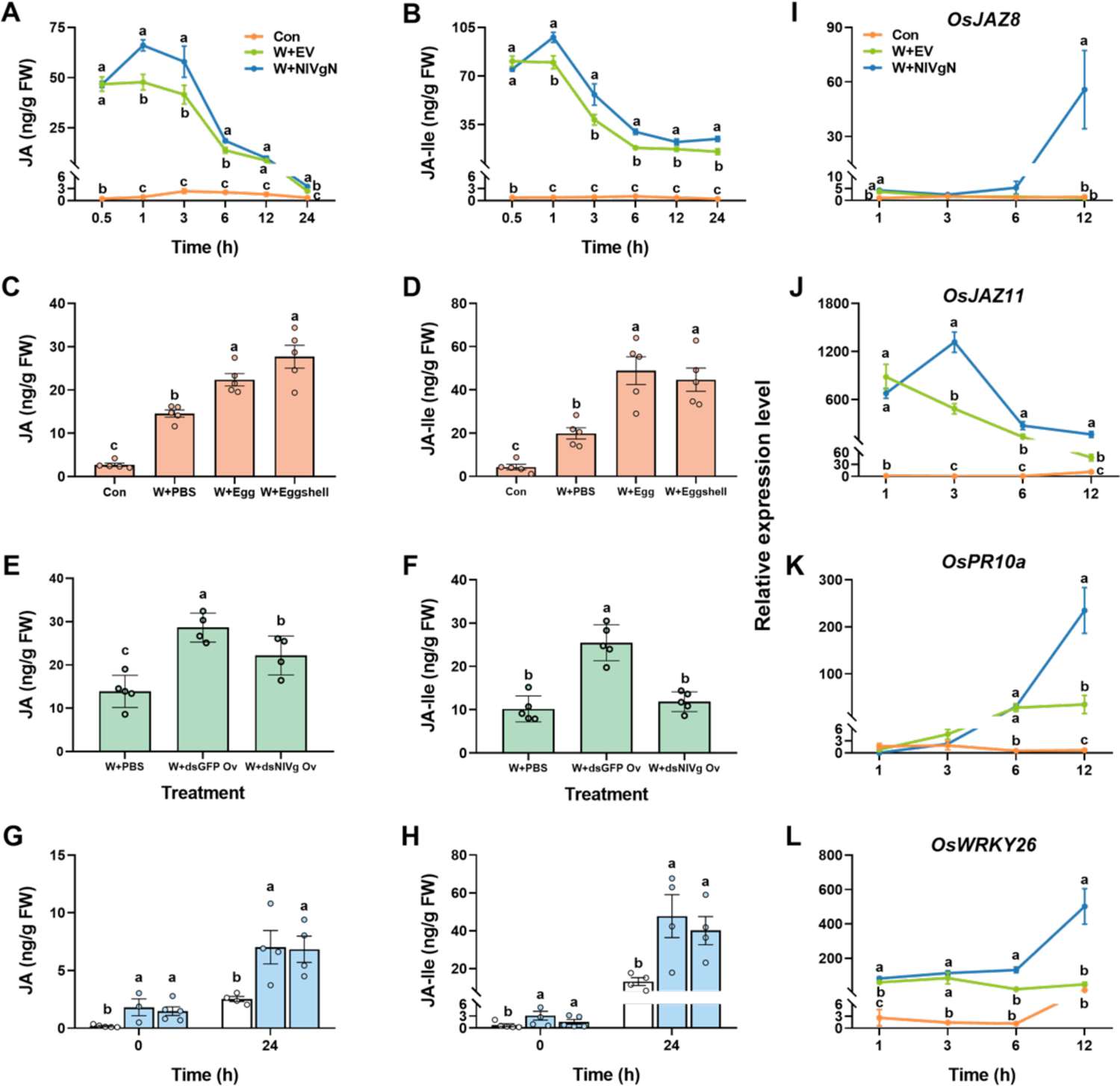
NlVgN elicits the production of JA and JA-Ile and the expression of defense-related genes in rice. **(A and B)** Mean levels (+SE, n = 5) of JA (**A**) and JA-Ile (**B**) in rice leaf sheaths that were kept unmanipulated (Con) or treated for 0.5, 1, 3, 6, 12 and 24 h with wounding plus the purified products of the empty vector (EV) (W+EV) or the purified recombinant protein NlVgN (W+NlVgN). (**C** and **D**) Mean levels (+SE, n = 5) of JA (**C**) and JA-Ile (**D**) in rice leaf sheaths that were kept unmanipulated (Con) or treated for 3 h with wounding plus the phosphate buffered saline (PBS) (W+PBS) or the solution of homogenized BPH eggs (W+Egg) or eggshells (W+Eggshell) in the buffer. (**E** and **F**) Mean levels (+SE, n = 5) of JA (**E**) and JA-Ile (**F**) in rice leaf sheaths that were treated for 3 h with wounding plus the PBS or the solution of homogenized ovaries of BPH female adults (4 d after emergence) that were injected with dsRNA of *GFP* (*dsGFP*) (W+dsGFP Ov) or *NlVg* (*dsNlVg*) (W+dsNlVg Ov) at fifth-instar nymph stage. (**G** and **H**) Mean levels (+SE, n = 5) of JA (**G**) and JA-Ile (**H**) in WT plants and plants expressing *NlVgN* (line oe-1 and oe-3) that were kept non-infested (0 h) or infested with 10 gravid BPH female adults for 24 h. (**I**-**L**) Mean transcript levels (+SE, n = 5) of *OsJAZ8* (**I**)*, OsJAZ11* (**J**), *OsPR10a* (**K**) and *OsWRKY26* (**L**), in rice leaf sheaths that were kept unmanipulated (Con) or treated for 1, 3, 6, and 12 with W+EV or W+NlVgN. FW, fresh weight. Letters indicate significant differences among different treatments (*P* < 0.05, Tukey’s HSD post-hoc test).

### NlVgN induces the expression of defense-related genes and defense response of rice to BPH infestation

Because NlVgN triggers the production of JA, JA-Ile and H_2_O_2_ in rice (Figure 2 and Figure 3) and because these molecule-mediated signaling pathways play an important role in modulating direct and indirect defenses of rice against BPH (Zhou et al., 2009; Xiao et al., 2012; Hu et al., 2016), we hypothesized that treating plants with recombination protein NlVgN will not only alter the expression of defense-related genes, but also affect the performance of BPH and the behavioral response of *Anagrus nilarpavatae*, an egg parasitoid of rice planthoppers. As predicted, the transcript level of three JA-responsive genes, *OsJAZ8* (Yamada et al., 2012; Xu et al., 2021), *OsJAZ11* (Xu et al., 2021), and *OsPR10a* (Ersong et al., 2021), and one defense-related gene, *OsWRKY26* (Li et al., 2021), in rice, were up-regulated after NlVgN treatment (Figure 3I-L).

The survival and the mass of BPH nymphs and newly-emerged BPH female adults fed on rice plants that were treated with wounding plus recombination protein NlVgN were similar to those fed on plants that were treated with wounding plus purified elution products of the empty vector (Figure 3—figure supplement 1D-F). In contrast, the hatching rate of BPH eggs and the number of eggs laid by 15 gravid BPH females (for 24 h) were significantly lower on plants that were twice treated with NlVgN than EV-treated plants. One-time treatment with NlVgN did not influence the hatching rate of BPH eggs and the number of eggs laid by 15 gravid BPH females (Figure 4A and B). Expressing *NlVgN* in plants also reduced the hatching rate of BPH eggs (Figure 4C and D).

**Figure 4.**
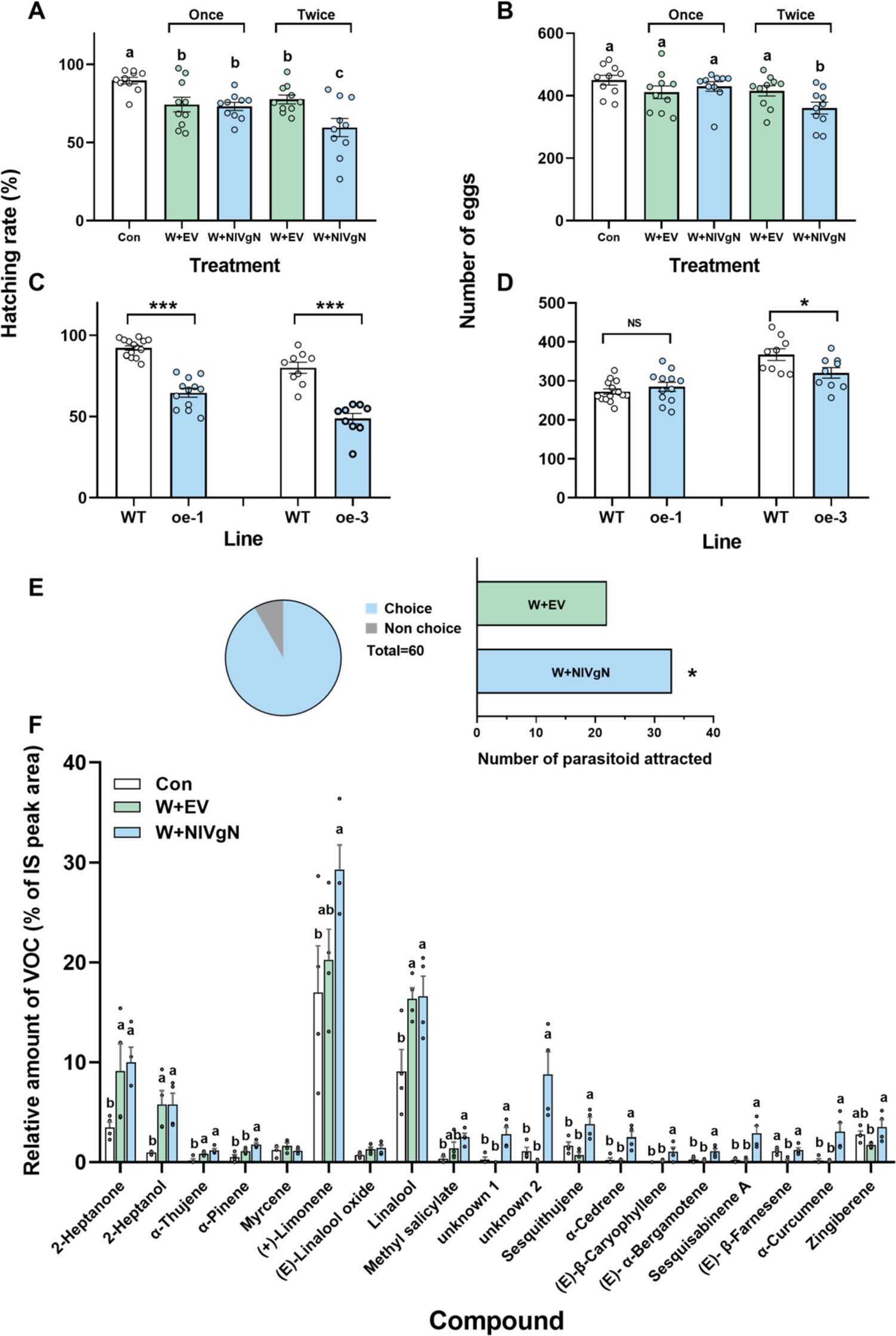
NlVgN induces direct and indirect defenses of rice against BPH. (**A-D**) Mean hatching rate (+SE, n = 9-15) of BPH eggs (**A** and **C**) and mean number (+SE, n = 9-15) of eggs laid by gravid BPH females for 24 h (**B** and **D**) on plants that were kept unmanipulated (Con), treated with wounding plus the purified products of the empty vector (EV) (W+EV) or the purified recombinant protein NlVgN (W+NlVgN), or expressed with *NlVgN* (line oe-1 and oe-3) or not (wild-type, WT). Once and twice indicate that plants were treated with W+EV or W+NlVgN one time and two times, respectively. The experiments on the hatching rate of BPH eggs on WT plants versus one of the two transgenic lines, oe-1 and oe-3, were performed separately. Asterisk indicates significant differences between different treatments (\**P* < 0.05; \*\*\**P* < 0.001; Student’s *t*-test). (**E**) Number of *A. nilaparvatae* female adults attracted by volatiles emitted from rice plants treated with W+EV or W+NlVgN. Asterisk indicates significant differences between different treatments (\**P* < 0.05, chi-squared test). (**F**) Mean amount (% of IS peak area, +SE, n = 4) of volatiles emitted from rice plants that were kept unmanipulated (Con) or treated with W+EV or W+NlVgN. Letters indicate significant differences among different treatments (*P* < 0.05, Tukey’s HSD post-hoc test).

Furthermore, volatiles emitted from plants treated with wounding plus purified recombination protein NlVgN were more attractive to female *A. nilaparvatae* wasps than volatiles from plants treated with wounding plus EV (Figure 4E). Volatile collections and analyses revealed that the total amount of volatiles emitted from plants treated with wounding plus purified recombination protein NlVgN was significantly higher than the total amount of volatiles from plants treated with wounding plus EV or from unmanipulated plants. Compared to non-manipulated control plants, wounding plus EV enhanced levels of four volatile compounds, 2-heptanone, 2-heptanol, α-thujene, and linalool, whereas nine volatile compounds, α-pinene, sesquithujene, α-cedrene, (*E*)-β-caryophyllene, (*E*)-α-bergamotene, sesquisabinene A, α-curcumene, and two unknown compounds, were released in higher amounts from recombination protein NlVgN-treated plants compared to control plants (Figure 4F). Taken together, the data imply that NlVgN induces the expression of defense-related genes and enhances the direct and indirect defense responses to BPH infestation.

### Silencing *NlVg* impairs BPH feeding, survival and fecundity

Consistent with results reported in Shen et al., 2019, knockdown of *NlVg* significantly increased the body size and mass of BPH female adults (Figure 5—figure supplement 1A-C), but resulted in oocyte malformations and drastically reduced the number of mature eggs in the ovaries (Figure 5—figure supplement 1D and 1E), as well as the number of eggs laid by female adults (Figure 5—figure supplement 1F). Knockdown of *NlVg* also decreased the amounts of honeydew secreted by newly-emerged female adults of BPH compared to those secreted by newly-emerged females of BPH injected with *dsGFP* (dsGFP-BPH) and BPH females that were not injected (C-BPH) (Figure 5A). Moreover, compared with dsGFP-BPH and C-BPH, dsNlVg-BPH showed lower survival rates on rice plants and artificial diet 6-10 d and 3-10 d, respectively, post injection (Figure 5B and C). The data confirm that NlVg plays an important role in the feeding, development, survival and especially fecundity of BPH.

**Figure 5.**
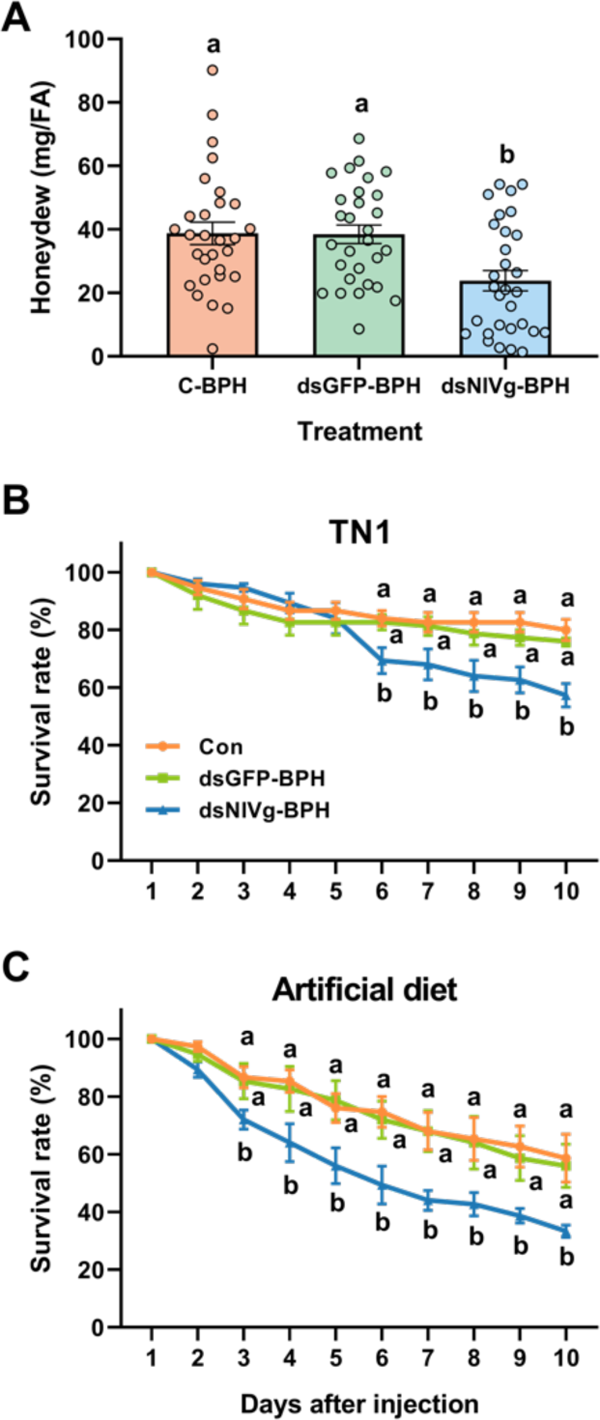
Knockdown of *NlVg* impairs the feeding capacity and survival of BPH female adults. (**A**) Mean amount of honeydew per day (+SE, n = 30) secreted by a newly-emerged BPH female adult (FA, 12-24 h after emergence) that was injected with dsRNA of *GFP* (*dsGFP*) or *NlVg* (*dsNlVg*), or kept non-injected (C-BPH) at fifth-instar nymph stage. (**B** and **C**) Mean survival rates (+SE, n = 5) of newly-emerged BPH female adults that were injected with *dsGFP*, *dsNlVg* or kept non-injected (C-BPH) at fifth-instar nymph stage, 1-10 d (2-11 d post injection) after they fed on rice variety TN1 (**B**) or artificial diet (**C**). Letters indicate significant differences among treatments (*P* < 0.05, Tukey’s HSD post-hoc test).

### VgNs in other rice planthoppers also function as elicitors

Rice plants suffer from attacks by several planthopper species. The main ones are BPH, WBPH and the small brown planthopper (SBPH) *Laodelphax striatellus*. We wondered whether VgNs in these rice planthoppers also induce defense responses in rice. We therefore investigated the change in levels of JA and JA-Ile in rice plants when they were treated with eggs or ovary solutions of WBPH or SBPH. Similar to results found for BPH, wounding plus applying the homogenized fresh egg or ovary solution of WBPH or SBPH resulted in higher levels of JA and JA-Ile than wounding plus applying the buffer (Figure 6A-D). When *Vg* was knocked down in WBPH or SBPH (Figure 6—figure supplement 1), the same treatments with WBPH or SBPH ovary solution did not or only weakly induce the production of JA-Ile (Figure 6C and D). Taken together, these findings show that VgN from WBPH and SBPH also functions as an elicitor that induces defense responses in rice.

**Figure 6.**
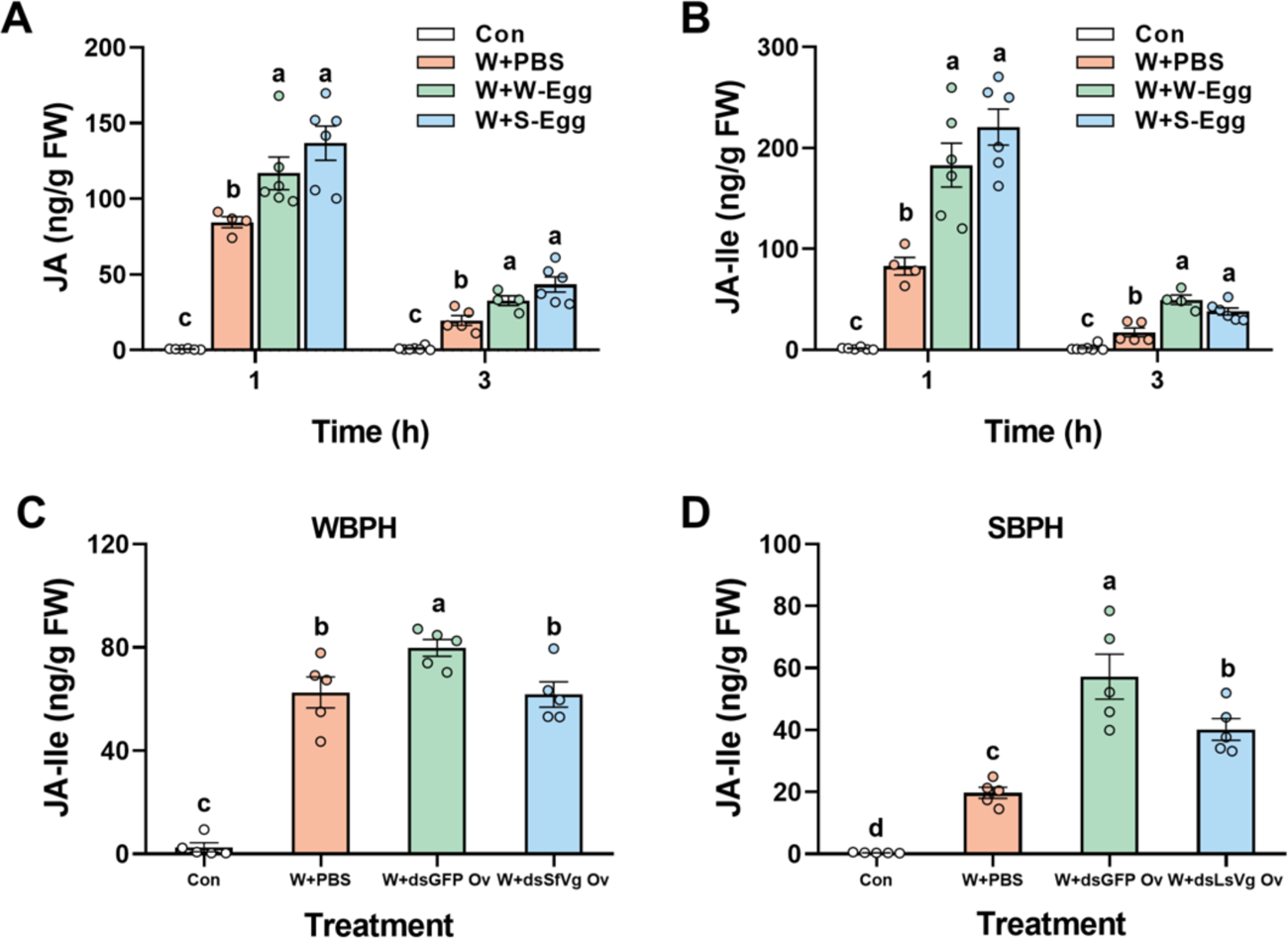
VgN in WBPH and SBPH also elicits the production of JA and JA-Ile. (**A** and **B**) Mean levels (+SE, n = 4-7) of JA (**A**) and JA-Ile (**B**) in rice leaf sheaths that were kept unmanipulated (Con) or treated with wounding plus the phosphate buffered saline (PBS) (W+PBS) or the solution of the homogenized WBPH eggs (W+W-Egg) or SBPH eggs (W+S-Egg) (for 1 and 3 h). (**C** and **D**) Mean levels (+SE, n = 5) of JA-Ile in rice leaf sheaths that were kept unmanipulated (Con) or treated for 1 h with W+PBS or the solution of homogenized ovaries of WBPH (**C**) or SBPH (**D**) female adults (4 d after emergence) that were injected with dsRNA of *GFP* (dsGFP) (W+dsGFP Ov), *SfVg* (dsSfVg) (W+dsSfVg Ov) or *LsVg* (dsLsVg) (W+dsLsVg Ov), respectively, at fifth-instar nymph stage. FW, fresh weight. Letters indicate significant differences among different treatments (*P* < 0.05, Tukey’s HSD post-hoc test).

## Discussion

The evolutionary arms race between plants and herbivorous insects has resulted in numerous clever defense traits in plants, and equally ingenious counter adaptations in specialized insects (Farmer, 2014). Plant defenses against insects are often inducible (Karban and Baldwin, 1997), and in order for the plants to launch the most appropriate defense they need to recognize their specific attackers. This is possible thanks to insect specific elicitors, also referred to as herbivore-associated molecular patterns (HAMPs) (Felton and Tumlinson, 2008; Arimura, 2021). In turn, the insect herbivores are under strong selective pressure to avoid excreting such indicative elicitors. This implies that only compounds that insects cannot avoid producing can serve as reliable elicitors. This is indeed the case for the two most studied types of HAMPs, fatty acid conjugates like volicitin (Hettenhausen et al., 2014; Yoshinaga et al., 2010) and inceptins, which are peptide fragments from chloroplastic ATP synthase γ-subunit proteins (Schmelz et al., 2006). Both types of elicitors are formed in caterpillar buccal cavities during feeding and cannot be avoided unless the insects adapt their diet (De Moraes and Mescher, 2004) or change digestive enzyme activity (Schmelz et al., 2012), respectively. Recently, the first plant receptor to allow this specific recognition of HAMPs was identified (Steinbrenner et al., 2020).

Here we identified a new type of elicitor, the small N-terminal subunit of vitellogenin protein from the brown planthopper (BPH), NlVgN. It is uniquely different from other elicitors in that it is introduced into the plants, not only via the saliva, but also, in large quantities, during oviposition. It activates different defensive signaling pathways and thereby causes various defense responses in rice. We also show that NlVgN is essential for BPH growth, development and fecundity. Hence, NlVgN is an unavoidable HAMP that highly reliable betrays the presence of BPH and other planthoppers. Below, we discuss the mode of action of NlVgN and elaborate on the evolutionary and possible pest control implications of our findings.

It is known that infestation by gravid BPH female adults (feeding + oviposition) activates cytosolic Ca^2+^ signaling and enhances levels of H_2_O_2_, JA, JA-Ile and SA, but it inhibits the production of ethylene, whereas infestation by BPH nymphs or newly-emerged female adults (feeding only; they do not lay eggs until after they go through a pre-oviposition period) only induces an increase in levels of cytosolic Ca^2+^, H_2_O_2_ and SA (Li, 2015; Ye et al., 2017). Based on our results we propose that the difference in responses to gravid BPH females and BPH nymphs is at least in part due to the difference in the source of NlVgN, in addition to different types of damage inflicted by nymphs and gravid females. NlVgN enters the plants at feeding sites via BPH saliva, as well as oviposition sites via the eggs (Figure 1E-I). The importance of this for defense induction was confirmed by the observation that knockdown of *NlVg* in the insect significantly decreased BPH feeding-induced levels of cytosolic Ca^2+^ and H_2_O_2_ (Figure 2A-D) but did not affect JA and JA-Ile levels (Figure 3—figure supplement 1A and 1B). Moreover, wounding plus the application of solutions with either homogenized fresh BPH eggs, eggshells or BPH ovaries, all of which contain NlVgN, induced the biosynthesis of JA and JA-Ile in plants (Figure 3C-F), whereas knocking down *NlVg* reduced the levels of JA and JA-Ile induced by the ovary solution (Figure 3E and F). Importantly, exogenous application of the recombinant NlVgN or expressing *NlVgN* in the rice plants themselves was sufficient to elicit the production of H_2_O_2_ (Figure 2E-G), JA and JA-Ile (Figure 3A and B, Figure 3G and H). These results imply that NlVgN from saliva induces the production of cytosolic Ca^2+^ and H_2_O_2_, whereas NlVgN from eggs induces the production of cytosolic Ca^2+^, H_2_O_2_, as well as JA and JA-Ile. This appears to be the first report on the role of vitellogenin in inducing plant defense responses, but, interestingly, vitellin produced by the cattle tick *Boophilus microplus* has been previously found to act as an elicitor of immune responses in sheep (Tellam et al., 2002). Recently, it was reported that the C-terminus of Vg, VgC, in SBPH, when secreted into rice plants, serves as an effector (Ji et al., 2021). This was concluded from the fact that *Vg*-silenced SBPH nymphs consistently elicited higher H_2_O_2_ production, whereas expression of the domains in VgC in rice protoplasts or of VgC in *Nicotiana benthamiana* leaves significantly hindered the accumulation of chitason-induced H_2_O_2_. Moreover, silencing *Vg* reduced SBPH feeding and survival on rice. The discrepancy between the conclusion from the SBPH study (Ji et al., 2021) and ours is possibly due to the fact that, besides the possibility that different peptides of VgN and VgC cleaved from Vg, in the SBPH study there was no calibration for the differences in damage levels caused by SBPH feeding on plants with different treatments. There is also no direct evidence that VgC indeed inhibits the production of SBPH-induced H_2_O_2_ in rice and improves the performance of SBPH on rice.

The reason why NlVgN from different sources differently affected the production of JA and JA-Ile might be related to the type and extent of damage caused and the compounds that enter into plants during BPH feeding versus oviposition. BPH is a piercing-sucking herbivore whose feeding only causes minor tissue damage and only little NlVgN will enter via the stylet sheaths. In contrast, during oviposition BPH causes considerably more damage to tissues as it needs to make cuts with its ovipositor to lay egg clusters inside the leaves (Figure 1B-D). As a consequence, NlVgN from the egg surface comes in direct contact with damaged tissues and thereby induces considerably stronger responses in rice plants than the small quantities of NlVgN deposited during feeding. Moreover, the fluids deposited by BPH also contain effectors and other elicitors, which are probably different between saliva and eggs/oviposition fluids. To date, one elicitor, a mucin-like protein (Shangguan et al., 2018), and several effectors, such as an endo-*β*-1,4-glucanase (Ji et al., 2017), an EF-hand calcium-binding protein (Ye et al., 2017), and 6 other proteins (Rao et al., 2019), from BPH saliva have been reported. The different combinations of these effectors and elicitors can explain why rice plants respond somewhat differently to NlVgN during feeding and oviposition.

Increases in levels of JA and JA-Ile are known to regulate the resistance of rice to BPH (Xu et al., 2021). We therefore investigated the effect of NlVgN treatment of rice plants on the performance of BPH on these plants. This revealed that when plants were treated with NlVgN twice, NlVgN-induced defenses decreased the hatching rate of BPH eggs and the number of eggs laid by BPH female adults, but when the plants were treated only once with NlVgN, there was no effect on any BPH performance parameter (Figure 4A and B; Figure 3—figure supplement 1D-F). The effect of NlVgN-induced defenses on the hatching rate was also observed in plants expressing *NlVgN* (Figure 4C and D). Future research will have to elucidate which defensive compounds cause the death of BPH eggs.

In rice, JA- and ethylene-mediated signaling pathways also regulate the biosynthesis of inducible volatiles (Lou and Cheng, 2005; Tong et al., 2012; Lu et al., 2014). NlVgN did not affect the production of ethylene in rice (Figure 3—figure supplement 1C). Hence, the fact that exogenous application of NlVgN increased the amounts of volatiles emitted from rice plants was probably due to NlVgN activation of JA and JA-Ile signaling. Compared to control plants, NlVgN-treated plants produced higher levels of 9 volatile compounds, all of which were also induced by gravid BPH female infestation (Xiao et al., 2012; Lu et al., 2014). However, the NlVgN-induced volatile blend was not exactly the same as the blend induced by gravid BPH females; 2-heptanone, linalool, limonene and methyl salicylate were induced by the latter but not by the former (Tong et al., 2012). Again, this discrepancy is probably due to effectors and other elicitors in BPH saliva and eggs/oviposition fluids. The changes in volatile emissions implies that NlVgN is not limited to direct defenses, but also involves volatile-mediated indirect defense that results in the attraction of parasitoids. Of the nine volatile compounds that showed increases after NlVgN treatment (Figure 4F), (*E*)-β-caryophyllene has been reported to be attractive to *A. nilaparvatae* (Lou and Cheng, 2005; Xiao et al., 2012). Therefore, the higher attractiveness of NlVgN-treated plants to this egg parasitoid (Figure 4E) was probably due to increases in this and possibly other volatiles.

We further show that the role of VgN in inducing rice defenses is not limited to BPH. Solutions with homogenized fresh eggs of two other planthoppers commonly found on rice, WBPH or SBPH, also induced the biosynthesis of JA and JA-Ile when applied to mechanically wounded plants (Figure 6A and B). Moreover, when *Vg* in WBPH or SBPH was knocked down, the induction of WBPH or SBPH ovary solution on the production of JA-Ile in mechanically wounded plants significantly decreased (Figure 6C and D). VgN is also found in the saliva and eggs of WBPH and SBPH (Huang et al., 2018) and the similarity of the amino acid sequence of VgN among the three planthoppers is high (84.38%; Figure 6—figure supplement 2). Taken together, these results indicate that the VgN derived from the three planthoppers functions as a common elicitor during the interaction between rice and planthoppers.

Vgs and their homologues have been reported to play an important role in the growth, development, survival and fecundity in many insect species, including BPH (Shen et al., 2019). We confirmed this by silencing *NlVg*, which caused abnormal increases in the body size and mass of BPH female adults and drastically decreased their fecundity and resulted in failed egg formation in the ovaries (Figure 5—figure supplement 1) (Shen et al., 2019). Moreover, silencing *NlVg* impaired BPH feeding and decreased its survival on rice plants and artificial diet (Figure 5). The increase in body size and mass of dsNlVg-BPH female adults is probably at least in part related to the failure of egg formation, which prevents them from laying eggs like normal female adults (laying eggs decreases their body size and mass). These findings demonstrate that NlVgN, like Vgs in other insects, is required for growth, development, survival and fecundity. Hence, the results not only support our hypothesis that VgNs play a key role in inducing defense responses in rice plants, they also reinforce the notion of evolutionary stability and that only compounds that are essential for insect performance and survival can be dependably exploited by plants as elicitors. By confirming this key importance of Vgs for planthoppers we also expose the vulnerability of *NlVg* and similar genes in other insects, making them excellent targets for gene silencing strategies to control pests (Christiaens et al., 2020). This is evident from our silencing experiment, which resulted in increased BPH mortality (Figure 5B and C) and almost completely impaired egg production (Figure 5—figure supplement 1D-F).

In summary, our study shows that VgN, the small N-terminal subunit of vitellogenin from rice planthoppers, readily enters rice tissues during planthopper feeding and oviposition. VgN from the saliva of rice planthoppers, together with the damage caused by planthopper feeding, possibly in combination with effectors and other elicitors, induces the production of cytosolic Ca^2+^ and H_2_O_2_, whereas VgN from eggs, accompanying the damage caused during oviposition and possible other chemical factors in oviposition fluids, induces the production of cytosolic Ca^2+^, H_2_O_2_, JA and JA-Ile. The activated JA signaling pathway decreases the hatching rate of BPH eggs and the number of eggs laid by BPH female adults, and increases the emission of volatiles from rice, which enhances the attractiveness of rice plants to the egg parasitoid *A. nilaparvatae*. Our study not only identifies VgN from rice planthoppers as a potent elicitor but also provides a compelling example of how an elicitor combined with varying herbivore inflicted damage types can cause differential defense responses in plants. The importance of VgNs for the planthoppers makes them stable and reliable indicators of planthopper presence and possible targets for molecular pest control strategies.

## Materials and methods

### Plant Growth

Rice genotypes used in this study were Taichun Native 1 (TN1), Xiushui 11 (wild type) and transgenic lines expressing *NlVgN* (oe-1 and oe-3; see details below); TN1 and Xiushui 11 are two rice varieties susceptible to BPH. Pre-germinated seeds were cultured in plastic bottles (diameter 8 cm; height 10 cm) in a greenhouse (27±1℃, 14/10 h light/dark photoperiod). Ten-day-old seedlings were transferred to 20-L hydroponic boxes with a rice nutrient solution (Yoshida et al., 1976), and 30- to 35-day-old plants were transferred to individual 500-mL hydroponic plastic pots for experiments. Plants used in all of experiments except for specified experiments were TN1 plants.

### Insects

A colony of BPH was originally provided by the Chinese National Rice Research Institute (Hangzhou, China) and maintained on TN1 plants in a climate chamber at 27±1 ℃ and 80% relative humidity under a 14/10 h light/dark photoperiod. Colonies of WBPH and SBPH were originally collected from rice fields in Hangzhou, China and maintained on TN1 plants in the climate chamber.

### Cloning of *NlVgN* and Sequence Analysis

The full-length cDNA of *NlVgN* was obtained by reverse transcription (RT)-PCR from total RNA isolated from gravid BPH females. Specific primers (Supplementary file 1) were designed based on the sequence of *NlVg* (GenBank: AB353856). PCR-amplified fragments were cloned into the *pEASY*^®^-Blunt Cloning Vector (Transgen, Beijing, China) and sequenced. DNAMAN (www.lynnon.com/) was used to deduce the amino acid sequences of *NlVgN* and to analyze the molecular weight of the predicted protein. Amino acid sequences of VgNs from WBPH (SfVgN) and SBPH (LsVgN) have been reported previously (Huo et al., 2018; Hu et al., 2019). The multiple sequence alignment analysis of VgNs from the three rice planthoppers was performed by the Clustal Omega (https://www.ebi.ac.uk/Tools/msa/clustalo/).

### Plant treatment

For mechanical wounding treatments, leaf sheaths of individual plant shoots (4 cm length) were punctured 80 times using a #3 insect pin. Unmanipulated plants were used as controls (Con). For NlVgN treatment, plants were individually wounded as stated above, and then were individually treated with 40 μL of the recombinant protein His-NlVgN (26.8 ng μL^-1^) or the purified products of the empty vector (EV), or kept unmanipulated (Con). For ovary, egg or eggshell solution treatments, plants were wounded as stated above, and then were individually treated with 40 μL of ovary, egg or eggshell solution. To prepare these solutions, ovaries (dissected from female adults of BPH, WBPH or SBPH 4 d after emergence and washed with PBS three times), eggs (collected from plants that had been oviposited on by gravid females of BPH, WBPH or SBPH for 24 h) and eggshells (collected from plants on which BPH nymphs had hatched within 24 h) were separately collected. All the collections were then homogenized in PBS containing 1 mM PMSF and the final concentration of ovary, egg and eggshell in the solution was one ovary, 20 eggs and 40 eggshells per 20 μL solution, respectively. Plants were wounded as described above and then plants were individually treated with 40 μL of the solution of ovaries (Ovary), eggs (Egg) or eggshells (Eggshell) or PBS (containing 1 mM PMSF), or kept unmanipulated (Con). For BPH treatment, individual plant shoots were confined in the glass cylinders into which 20 (dsNlVg-BPH) or 15 (dsGFP-BPH and C-BPH) newly-emerged BPH female adults were introduced; the number of BPH with different treatments, dsNlVg-BPH, dsGFP-BPH or C-BPH, on each plant was determined according to the difference in their food intakes on plants (Figure 5A; ensuring that each plant received equal damage from BPH). To ensure there were no eggs laid on the plants by these BPH females, they were replaced, every 48h, with a new set of newly-emerged females with the same treatment. Plants with empty glass cylinders (Non-infested) and kept unmanipulated (Con) were used as controls. When plants expressing *NlVgN* and wild type (Xiushui 11) plants were used, each cylinder received 10 gravid C-BPH females for BPH treatments.

### RNA Extraction and Quantitative Real-time PCR Analysis

Total RNA was extracted from the following materials: (1) whole bodies of female adults of BPH, WBPH, or SBPH at 1, 3 and 5 d after emergence; (2) leaf sheaths of rice plants expressing *NlVgN* (lines oe-1 and oe-3) and wild type plants; (3) leaf sheaths of rice plants that were kept unmanipulated (Con) or had been treated for 1, 3, 6 and 12 h with wounding plus the purified recombinant protein NlVgN (NlVgN) or the purified products of the empty vector (EV). Total RNA was isolated using the SV Total RNA Isolation System (Promega Corporation, Madison, WI, USA) by following the manufacturer’s protocol. cDNA was synthesized from 500 ng of total RNA in a 10 μL reaction using the Takara Primescript™ RT reagent kit. qRT-PCR was performed with the CFX96^TM^ Real-Time system (Bio-Rad, Hecrules, CA, USA) using the SYBR Premix EX Taq Kit (Takara Bio Inc., Kusatsu, Japan). A relative quantitative method (2^-ΔΔCt^) described previously (Pfaffl, 2001) was applied to evaluate the variation in expression levels of target genes among samples. The expression level of target genes in planthoppers and rice plants was normalized to *RPS15* (BPH ribosomal protein S15e gene, for developmental stage) (Yuan et al., 2014), *RPL9* (WBPH ribosomal protein L9 gene, for WBPH) and *EF2* (SBPH elongation factor 2 gene, for SBPH), and *OsActin1* (for rice plants), respectively. The primers used for qRT-PCR analysis in this study were provided in Supplementary file 2. Three or five independent biological replicates were used.

### RNA Interference

A unique region of *NlVg*, *SfVg*, *LsVg* and *GFP* were amplified by PCR with primers containing the T7 promoter sequence at both ends (Supplementary file 1). The purified PCR products were used to synthesize dsRNA by using the MEGAscript T7 High Yield Transcription Kit (Ambion, Austin, TX, USA) according to the manufacturer’s instructions. The concentration of dsRNA was quantified and the quality of dsRNA was further verified via electrophoresis in a 1% agarose gel. Fifth-instar nymphs were injected using the same method as described previously (Liu et al., 2010). Each nymph was injected about 0.4 μg dsRNA of *Vg* or *GFP*, or kept unmanipulated. To detect the silencing efficiency, the transcript level of *Vg* in the whole bodies of BPH, WBPH or SBPH female adults that had been injected with dsRNA of *Vg* or *GFP*, or kept non-injected were determined at 1, 3 and 5 d after emergence (2, 4, and 6 d post injection). Three independent biological replicates were used.

### Expression of NlVgN in *Escherichia coli*

The open reading frame of *NlVgN* was amplified by PCR using a pair of primers listed in Supplementary file 1. The PCR product was cloned into the pET-28a vector (Novagen, Inc., Madison, WI, USA) and sequenced. The recombinant vector *NlVgN*:pET-28a (Supplementary file 3) and empty vector pET-28a (as a control) were transformed into *E. coil* BL21 (DE3) strain. The protein fused with His-tag was expressed after induction with 1 mM isopropyl β-D-1-thiogalactopyranoside (IPTG) at 16℃ for 12 h and purified by using Ni-NTA resin columns (Qiagen, Venlo, Netherlands) according to the manufacturer’s instructions. All products purified from recombinant vector or empty vector were mixed with 5×protein loading buffer, separated by SDS-PAGE (sodium dodecyl sulfate polyacrylamide gel electrophoresis) in a 12% (w/v) acrylamide gel, and stained with 0.025% Coomassia Blue R-250 in water. The predicted mass of the mature recombinant protein NlVgN containing six N-terminal His-tags is 52.46 kDa.

### Polyclonal Antibody Preparation and Western Blot Analysis

A polypeptide of NlVgN, NPASNSESNQRSSH was selected as the antigen to produce the polyclonal rabbit antibodies, and the polyclonal antibodies were purified by GenScript (Nanjing, China). Protein samples used for western blot analysis were prepared as follows: (1) Proteins from rice leaf sheaths infested by BPH or not. Individual plant shoots (0-8 cm above the ground) of TN1 were confined within a glass cylinder (diameter 4 cm, height 8 cm, with 48 small holes, diameter 0.8 mm) in which 50 newly-emerged BPH female adults (12-24 h after emergence) that were injected with dsRNA of *GFP* or *NlVg* were released, and 24 h later, the herbivores were removed. Plants in empty glass cylinders were used as controls. The outer two leaf sheaths for each plant were harvested and the entire leaf sheaths from three plants were merged and ground in liquid nitrogen; approximate 150 mg of samples were homogenized in 500 μL RIPA Lysis Buffer (Beyotime, Shanghai, China) containing 1 mM PMSF (Phenylmethanesulfonyl fluoride), and the extract was centrifuged at 12,000×g for 5 min at 4℃. The supernatant was collected and the protein concentration in the supernatant was measured by using Pierce BCA Protein Aaasy Kit (Thermo Fisher Scientific, Rockford, IL, USA). (2) Proteins from the surface of BPH eggs, as well as from pieces of rice leaf sheaths from which BPH eggs had been gently removed. For this, plants were individually infested with 20 gravid BPH females for 24 h. Two hundred intact eggs were carefully collected from rice leaf sheaths. Then the eggs and the pieces of leaf sheaths with eggs removed were separately placed into 40 μL PBS buffer in a centrifuge tube for 2-3 min and 8-10 min, respectively. The supernatant was collected. The supernatants of extracts from pieces of leaf sheaths that were infested by newly-emerged BPH females (no eggs) or kept unmanipulated were used as controls. As an additional control we used the PBS buffer alone. (3) Proteins extracted from the whole bodies of 1-, 3- and 5-day-old BPH female adults on the 2-, 4- and 6-days post injection with dsRNA of *NlVg* or *GFP* (as a control). Two BPH female adults were homogenized in 1 mL of phosphate-buffered saline (PBS, pH 7.4) containing 1 mM PMSF, and the extract was centrifuged at 12,000×g for 10 min at 4℃. The supernatant was collected and the protein concentration in the supernatant was measured. Protein samples (20 μg from plant samples, 3 μg from BPH samples, 20 μL from the solutions used to extract BPH eggs or pieces of leaf sheaths) (mixed with 5×protein loading buffer) were subjected to SDS-PAGE on a 12% gel and transferred onto a nitrocellulose membrane. The membrane was blocked overnight at 4℃ with skim milk (5%) in 1×TBST (Tris-buffered saline with 0.05% Tween-20), and then was washed with 1×TBST and incubated with anti-NlVgN antibody (1:5000) or anti-β-Actin (1:5000; Engibody, Dover, DE, USA) for 1 h at 37℃, followed by extensive washing for 30 min with frequent changes of 1×TBST. After this, the membrane was incubated with HRP-conjugated goat anti-rabbit antibodies (1:10000) or goat anti-mouse antiantibodies (1:5000) for 1 h at 37℃, followed by extensive washing for 20 min with frequent changes of 1×TBST. Western blots were imaged using the Clarity^TM^ Western ECL Substrate (Bio-Rad, Hercules, CA, USA) with the Molecular Imager^®^ ChemiDoc^TM^ XRS+ System (Bio-Rad, Hercules, CA, USA).

### Immunofluorescence Microscopy

To locate the position of NlVgN in rice tissues that were infested by gravid BPH females and in BPH tissues, the following materials were collected. (1) The outermost leaf sheath of rice plants that had been infested by gravid BPH females for 6 h; (2) Fresh BPH eggs dissected from leaf sheaths of rice plants that had been infested by gravid BPH females for 12 h. The specimens were fixed in 4% paraformaldehyde at room temperature for 2 h and three times washed in PBS. The specimens were then incubated in PBST/BSA (PBS containing 2% Tween-20 and 2% bovine serum albumin) at room temperature for 2 h, followed by incubated at room temperature for 1 h each with anti-NlVgN antibody (1:200 diluted in PBST/BSA) and Alexa 568-labeled goat anti-rabbit antibody (1:200 diluted in PBST/BSA, Invitrogen, Carlsbad, CA, USA). The specimens were washed three times in PBST then examined using a Zeiss LSM 800 confocal laser scanning microscope.

### Generation of Transgenic Plants

The full-length coding sequence of *NlVgN* was PCR-amplified using a pair of primers listed in Supplemental Table 1 and was digested by *Kpn*I and *Xba*I; the product was then cloned into the binary vector pCAMBIA1301, yielding an overexpression transformation vector *NlVgN*:pCAMBIA1301 (Supplementary file 3). The vector was inserted into the Xiushui 11 plants via *Agrobacterium tumefaciens*-mediated transformation. Rice transformation, screening of the transgenic plants and identification of the number of insertions were performed following the same method as described previously (Zhou et al., 2009). Two *NlVgN*-expressing lines at T_2_ generation, oe-1 and oe-3, each with one insertion (Figure 2—figure supplement 2), were used for experiments.

### BPH Bioassays

To investigate the effect of NlVgN on BPH feeding capacity, a newly-emerged brachypterous female adult of dsNlVg-BPH, dsGFP-BPH or C-BPH (1 d after the injection of dsRNA of *NlVg* or *GFP*, or no injection, respectively, at fifth-instar nymph stage) was introduced into a Parafilm bag (6×5 cm), which was then fixed onto the shoot of a rice plant. Twenty-four hours later, the amount of honeydew excreted onto the Parafilm by a female adult was weighed (to an accuracy of 0.1 mg; Sartorius, BSA124S-CW). Each treatment was replicated thirty times.

To determine the influence of NlVgN on the fecundity of BPH, a newly-emerged BPH female adult of dsNlVg-BPH, dsGFP-BPH or C-BPH as stated above and a newly-emerged male adult (non-injected) were confined in the shoot (0-8 cm above the ground) of a plant with the glass cylinder. Ten days later, the number of eggs laid by each female adult was counted under a microscopy. Each treatment was replicated eleven times.

To assess the effect of NlVgN on the growth of BPH, the mass of female adults of dsNlVg-BPH, dsGFP-BPH or C-BPH (to an accuracy of 0.1 mg) were weighed at 1, 3, 5, 7 and 9 d after emergence. Each treatment at each time point was replicated three to nine times, and each replication contained at least three BPHs.

The effect of NlVgN on the survival of BPH reared on rice plants or artificial diet was also investigated. Briefly, for the survival experiment on rice plants, individual plant shoots were confined within the glass cylinders and then 15 newly-emerged female adults of dsNlVg-BPH, dsGFP-BPH or C-BPH as stated above were introduced into each cylinder. For the survival experiment on AD, one open end of the glass cylinder was covered with two layers of stretched Parafilm membrane, which contained an artificial diet (Fu et al., 2001), then, fifteen newly-emerged female adults of dsNlVg-BPH, dsGFP-BPH or C-BPH were released as described above. The artificial diet was replaced every day. The number of surviving BPHs was recorded for 10 days. Five independent replications were performed.

To explore the effect of treatment with the recombination protein NlVgN on the survival of BPH nymphs or newly-emerged female adults, plants were randomly assigned to NlVgN, EV and control treatments. Twenty-four h later, individual plant shoots (0-8 cm above the ground) were confined within the glass cylinders into which 20 nymphs or 15 newly-emerged female adults of BPH were introduced. The number of the surviving BPHs was recorded every day for 10 days, and the mass of all BPHs was weighed at the end of the experiment. Five independent replications were performed.

To investigate the effect of treatment with the recombination protein NlVgN on the hatching rate of BPH eggs, plants were randomly assigned to NlVgN, EV and control treatments. These plants were divided into two groups each with NlVgN, EV and control treatments. Twelve h later, one of the two group of plants was retreated again (receiving the same treatments as before). Twenty-four h after the first treatment, fifteen gravid BPH females were allowed to oviposit on each plant of the two groups for 24 h. The number of newly-hatched nymphs on each plant was counted every day until no newborn nymph appeared for three consecutive days. Unhatched eggs in each plant were counted under the microscope to calculate the hatching rate of eggs. Ten replications for each treatment were performed.

We also measured the effect of plants expressing NlVgN on the hatching rate of BPH eggs. Plants of WT (Xiushui 11) and two transgenic lines were individually exposed to 15 gravid BPH females that were allowed to oviposit for 24 h. Then, the hatching rate of BPH eggs on WT and transgenic plants was calculated using the same method as above. Each line was replicated at least nine times.

### Intracellular Calcium Ion Variation Determination

Fluo-3 AM (acetoxy-methyl ester of Fluo-3) was used to determine the intracellular calcium ion variation as previously described (Ye et al., 2017). Briefly, a working solution of 5 μM Fluo-3 AM (stock solution in dimethyl sulfoxide) containing 50 mM MES (2-(N-morpholino) ethanesulfonic acid) buffer (pH 6.0), 0.5 mM calcium sulfate, 2.5 μM 3-(3,4-dichlorophenyl)-1,1-dimethylurea and 1% methanol. The shoots of TN1 plants were individually confined in the glass cylinders into which 15 newly-emerged female adults of dsNlVg-BPH or dsGFP-BPH were released as explained above. Infested parts of leaf sheaths (about 3 cm) were individually harvested 1 and 3 h after infestation, and were immediately incubated in 1 mL of 5 μM Fluo-3 AM working solution at 37℃ for 30 min. The samples were mounted on a Zeiss LSM 780 confocal laser scanning microscope and were observed at 488 nm excitation wavelength. Images generated by the Zen 2010 software were analyzed by using the ImageJ software (https://imagej.nih.gov/ij). The fluorescence intensity at BPH feeding sites was individually measured at least sixteen times.

### H_2_O_2_ Analysis

TN1 plants were randomly assigned to NlVgN, EV and control treatments; Transgenic and WT (Xiushui 11) plants were randomly assigned to BPH and control treatments. For NlVgN, EV and control treatment, leaf sheaths of each plant were harvested at 0.5, 1, and 3 h after the start of treatment. For BPH and control treatment, leaf sheaths of each plant were harvested at 3, 8, and 24 h after BPH infestation. H_2_O_2_ was extracted using the same method as described previously (Lou and Baldwin, 2006) and the concentration of H_2_O_2_ were determined using Amplex-Red Hydrogen Peroxide/Peroxidase Assay Kit (Invitrogen, Carlsbad, CA, USA), following the manufacturer’s instructions. Each treatment at each time point was replicated five times. We also analyzed H_2_O_2_ by in situ detection. Rice leaves were pierced and treated with 20 μL of the recombinant protein NlVgN (26.8 ng μL^-1^) or the purified products of the empty vector (EV), or kept unmanipulated (Con). Fifteen min later at room temperature, leaves were stained with 3,3’-Diaminobenzidine (DAB) as described previously (Asano et al., 2012).

### JA and JA-Ile Analysis

TN1 plants were randomly assigned to BPH, NlVgN, EV, Ovary, Egg, Eggshell, PBS and control treatments; WT (Xiushui 11) and transgenic plants were randomly assigned to BPH and control. For NlVgN, EV and control treatment, leaf sheaths were harvested at 0.5, 1, 3, 6, 12, and 24 h after treatment. For Ovary, Egg, Eggshell, PBS and control treatment, leaf sheaths were harvested at 1 and/or 3 h after treatment. For BPH treatments, leaf sheaths of TN1 plants were harvested at 48, 72, and 96 h after infestation by newly-emerged BPH non-ovipositing female adults; leaf sheaths of WT and transgenic plants were harvested at 0 and 24 h after gravid BPH female infestation. JA and JA-Ile were extracted with ethyl acetate spiked with labeled internal standards (^2^D_6_-JA and ^2^D_6_-JA-Ile) following the method described previously (Lu et al., 2015) and analyzed by HPLC-MS/MS. Each treatment at each time point was replicated at least five times.

### Ethylene Analysis

TN1 plants were randomly assigned to NlVgN, EV and control treatments. Ethylene accumulation from individual plants with different treatments was measured by GC at 24, 48, and 72 h after the start of the treatment using the same method as described previously (Lu et al., 2006). Each treatment at each time interval was replicated eight times.

### Volatile Collection and Isolation

TN1 plants were randomly assigned to NlVgN, EV and control treatments. Twelve h after treatment, the volatiles emitted from individual plants were collected (for 8 h), isolated and identified using the method described previously (Lou and Cheng, 2005). The compounds were expressed as a percentage of peak areas relative to the internal standard (IS, diethyl sebacate) per 8 h of trapping for one plant. Collections were replicated four times for each treatment.

### Olfactometer Bioassays

Behavioral responses of *A. nilaparvatae* females to rice volatiles were performed in a Y-tube olfactometer using the same method as described previously (Lou and Cheng, 2005). The attraction of the parasitoid females exposed to the following pair of odor sources was recorded: TN1 plants treated with wounding plus purified recombination protein NlVgN for 12 h versus TN1 plants treated with wounding plus purified elution products of the empty vector for 12 h. For each treatment, 8 plants were used, and the odor sources were replaced by a new set of 8 plants after testing 20 wasps. In total, three sets of plants and 60 female parasitoids were used.

## Data Analysis

Two-treatment data were analyzed using Student’s *t* tests or chi-squared test (olfactometer bioassays). Data from three or more treatment groups was compared using one-way ANOVA; if the ANOVA was significant (*P* < 0.05), Tukey’s honest significant difference (HSD) post-hoc test was used to detect significant differences between treatments. All statistical analyses were performed using IBM SPSS Statistics 20.

## Data availability

All data generated or analyzed during this study are included in the manuscript and supporting files. Source data files have been provided for all figures and figure supplements.

## Author contributions

J.Z., W.Y., T.C.J.T. and Y.L. conceived and designed the experiments. J.Z., W.Y., W.H., X.J., P.K., W.X., and Y.J. performed the experiments. J.Z., W.Y, W.H. and Y.L. analyzed the data. J.Z., T.C.J.T. and Y.L. wrote the manuscript. All authors have read and approved the final manuscript.

## Acknowledgments

Authors declare that they have no competing interests.

---

**Figure 2—figure supplement 1.**
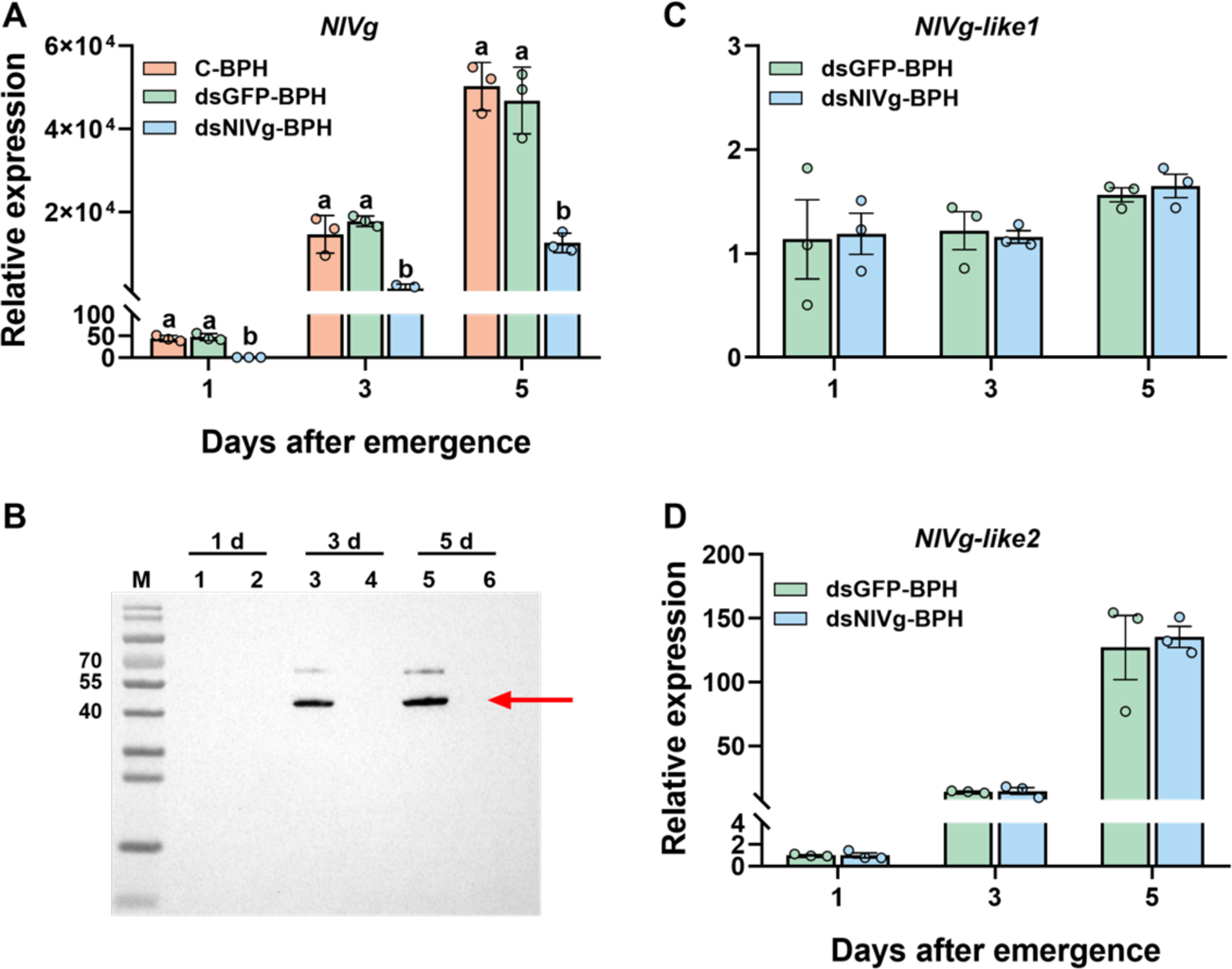
Silencing efficiency of *NlVg* by RNAi, and the effect of knocking down of *NlVg* on transcript levels of *NlVg-like1* and *NlVg-like2*. (A) Mean transcript levels (+SE, n = 3) of *NlVg* in whole bodies of 1-, 3- and 5-day-old BPH female adults 2, 4 and 6 d, respectively, after they were injected with dsRNA of *GFP* (*dsGFP*) or *NlVg* (*dsNlVg*), or kept non-injected (C-BPH) at fifth-instar nymph stage. Letters indicate significant differences among different treatments (*P* < 0.05, Tukey’s HSD post-hoc test). (B) Western blot analysis of NlVgN in proteins extracted from whole bodies of 1-, 3- and 5-day-old BPH female adults 2, 4 and 6 d, respectively, after they were injected with *dsGFP* (lanes 1, 3 and 5) or *dsNlVg* (lanes 2, 4 and 6) at fifth-instar nymph stage. M, molecular weight markers (kDa). (**C** and **D**) Mean transcript levels (+SE, n = 3) of *NlVg-like1* (**C**) and *NlVg-like2* (**D**) in whole bodies of 1-, 3- and 5-day-old BPH female adults 2, 4 and 6 d, respectively, after they were injected with *dsGFP* or *dsNlVg* at fifth-instar nymph stage. Differences between treatments at each time point are not significant (*P* > 0.05).

**Figure 2—figure supplement 2.**
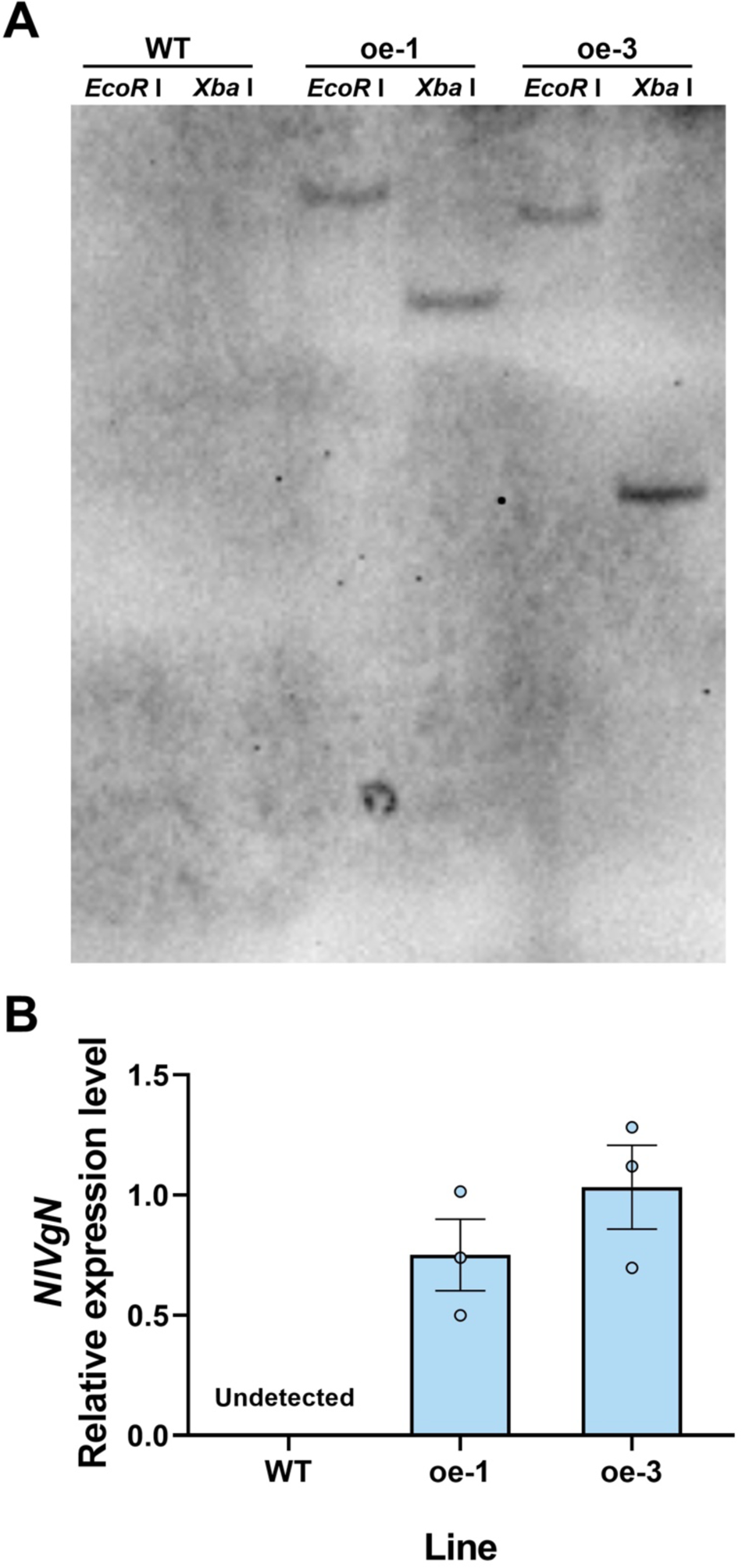
DNA gel-blot analysis and expression levels of *NlVgN* in transgenic (oe-1, oe-3) and wild-type (WT) plants. (**A**) Genomic DNA was digested with EcoRI and XbaI. The blot was hybridized with a probe specific for reporter gene *GUS* as *GUS* was inserted into the plant genome together with the target gene. The DNA-hybridized band of oe-1 and oe-3 means a single insertion by southern blotting. (**B**) Mean transcript levels (+SE, n = 3) of *NlVgN* in oe-1, oe-3, and WT plants that were kept unmanipulated.

**Figure 3—figure supplement 1.**
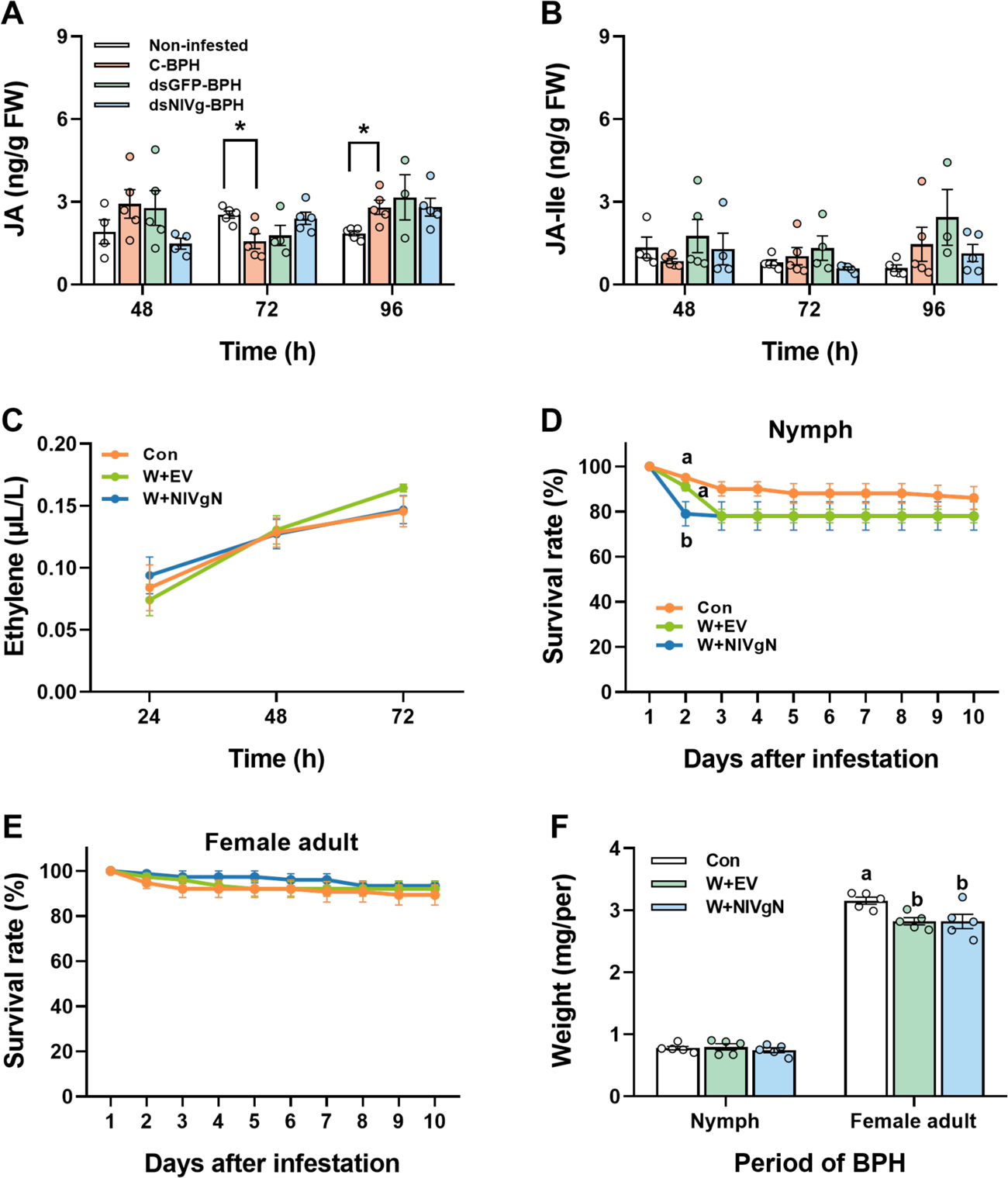
Effects of NlVgN secreted from BPH feeding or recombinant NlVgN protein on the production of JA, JA-Ile, or ethylene in rice, and on the survival and mass of BPH. (**A** and **B**) Mean levels (+SE, n = 5) of JA (**A**) and JA-Ile (**B**) in rice leaf sheaths that were kept insect-free (Non-infested) or individually infested by newly emerged BPH female adults which were injected with dsRNA of *GFP* (*dsGFP*), *NlVg* (*dsNlVg*) or kept noninjected (C-BPH) for 48, 72 and 96 h. Differences in the level of JA and JA-Ile between treatments at each time point are not significant (ANOVA, *P* > 0.05). Asterisks indicate significant differences between two treatments (\**P* < 0.05, Student’s *t*-test). (**C**) Mean levels (±SE, n = 8) of ethylene emitted from rice plants that were kept unmanipulated (Con) or treated with wounding plus the purified products of the empty vector (EV) (W+EV) or the purified recombinant protein NlVgN (W+NlVgN). Differences between treatments at each time point are not significant (ANOVA, *P* > 0.05). (**D** and **E**) Mean survival rates (±SE, n = 5) of newly hatched BPH nymphs (**D**) and of newly emerged female adults (**E**) 1-10 d after feeding on rice plants that were kept unmanipulated (Con) or treated with W+EV or W+NlVgN. (**F**) Mean mass (+SE, n = 5) of BPHs 10 d after feeding on rice plants that were kept unmanipulated (Con) or treated with W+EV or W+NlVgN. Letters indicate significant differences among different treatments (*P* < 0.05, Tukey’s HSD post-hoc test).

**Figure 5—figure supplement 1.**
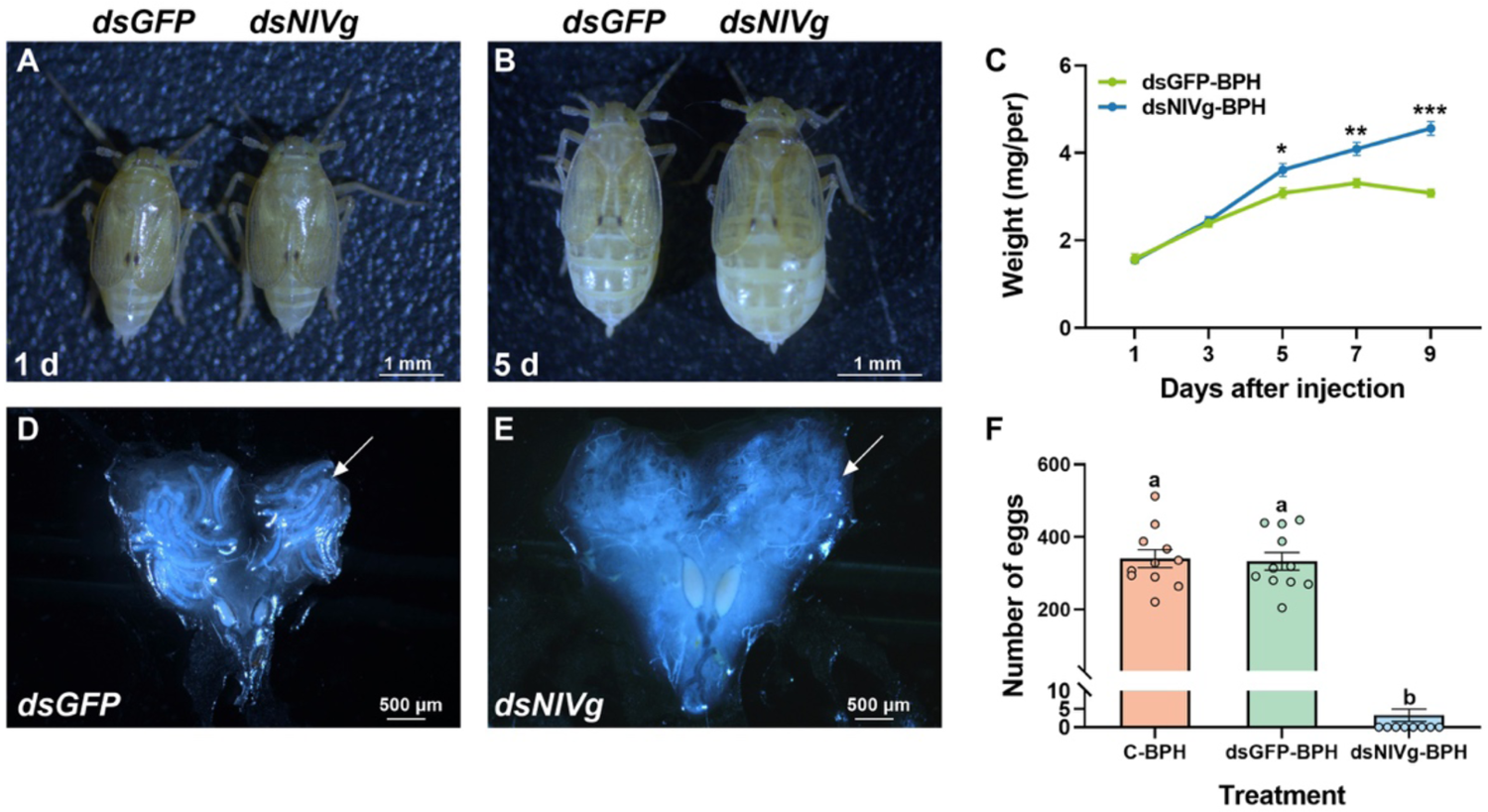
Knockdown of *NlVg* impairs the development and fecundity of BPH female adult. (**A** and **B**) The growth phenotypes of 1- (**A**) and 5-d-old female adults (**B**) at 2 and 6 d after they were injected with dsRNA of *GFP* (*dsGFP*) or *NlVg* (*dsNlVg*) at fifth-instar nymph stage. (**C**) Mean mass (±SE, n = 3-9) of individual 1-, 3-, 5-, 7- and 9-d-old female BPH adults at 2, 4, 6, 8 and 10 d after they were injected with *dsGFP* or *dsNlVg* at fifth-instar nymph stage. Asterisks indicate significant differences between different treatments (\**P* < 0.05, \*\**P* < 0.01, \*\*\**P* < 0.001, Student’s *t*-test). (**D** and **E**) The ovarian phenotypes of 5-d-old-female adults at 6 d after they were injected with *dsGFP* (**D**) or *dsNlVg* (**E**) at fifth-instar nymph stage. (**F**) Mean number of eggs (+SE, n = 11) on rice plants laid for 10 d by a female adult that was injected with *dsGFP* or *dsNlVg*, or kept non-injected (C-BPH) at fifth-instar nymph stage. Letters indicate significant differences among treatments (*P* < 0.05, Tukey’s HSD post-hoc test).

**Figure 6—figure supplement 1.**
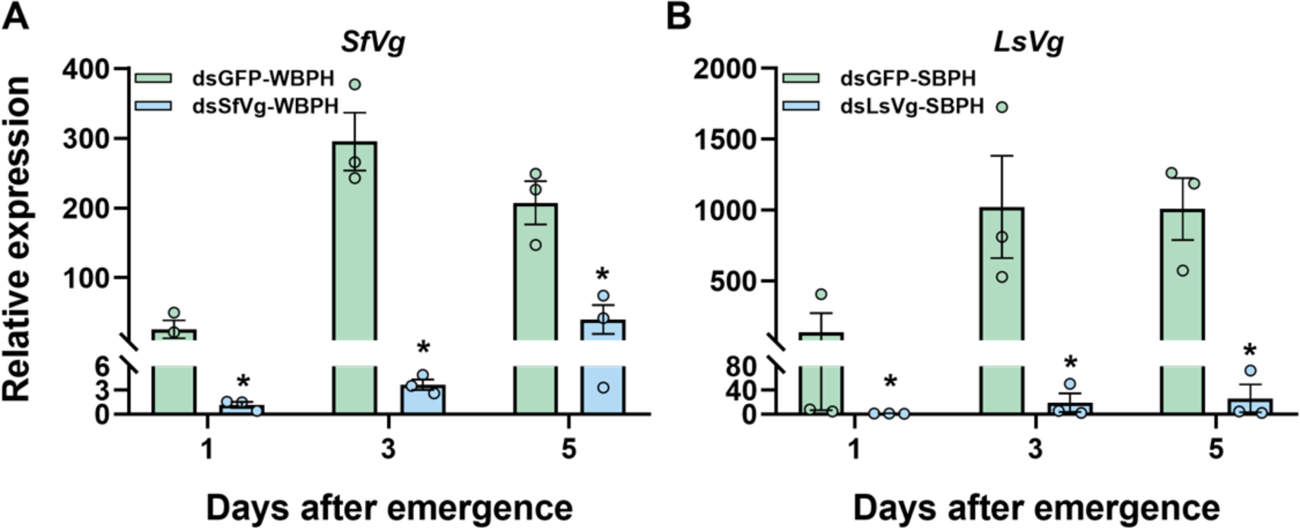
Silencing efficiency of *SfVg* and *LsVg* by RNAi in transcript levels. Mean transcript levels (+SE, n = 3) of *SfVg* (**A**) and *LsVg* (**B**) in whole bodies of 1-, 3- and 5-d-old female adults 2, 4 and 6 d after they were injected with dsRNA of *GFP* (*dsGFP*), S*fVg* (*dsSfVg*) or *LsVg* (*dsLsVg*) at fifth-instar nymph stage. Asterisks indicate significant differences between different treatments (\**P* < 0.05, Student’s *t*-test).

**Figure 6—figure supplement 2.**
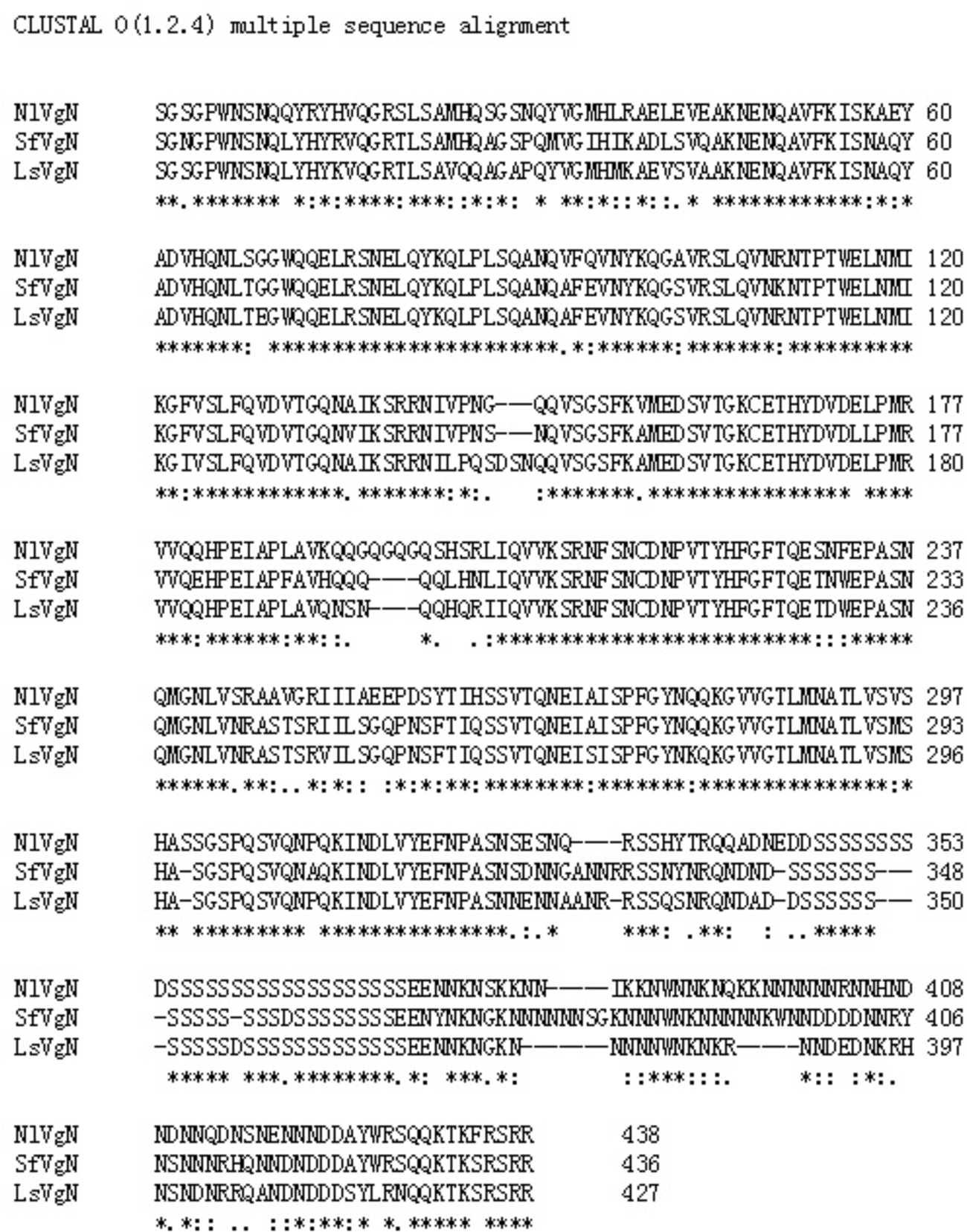
Multiple alignment of N-terminal subunit amino acid sequences of Vgs of three planthoppers. *, identical amino acid; :, conserved substitution; ., semiconserved substitution.

**Supplementary file 3.**
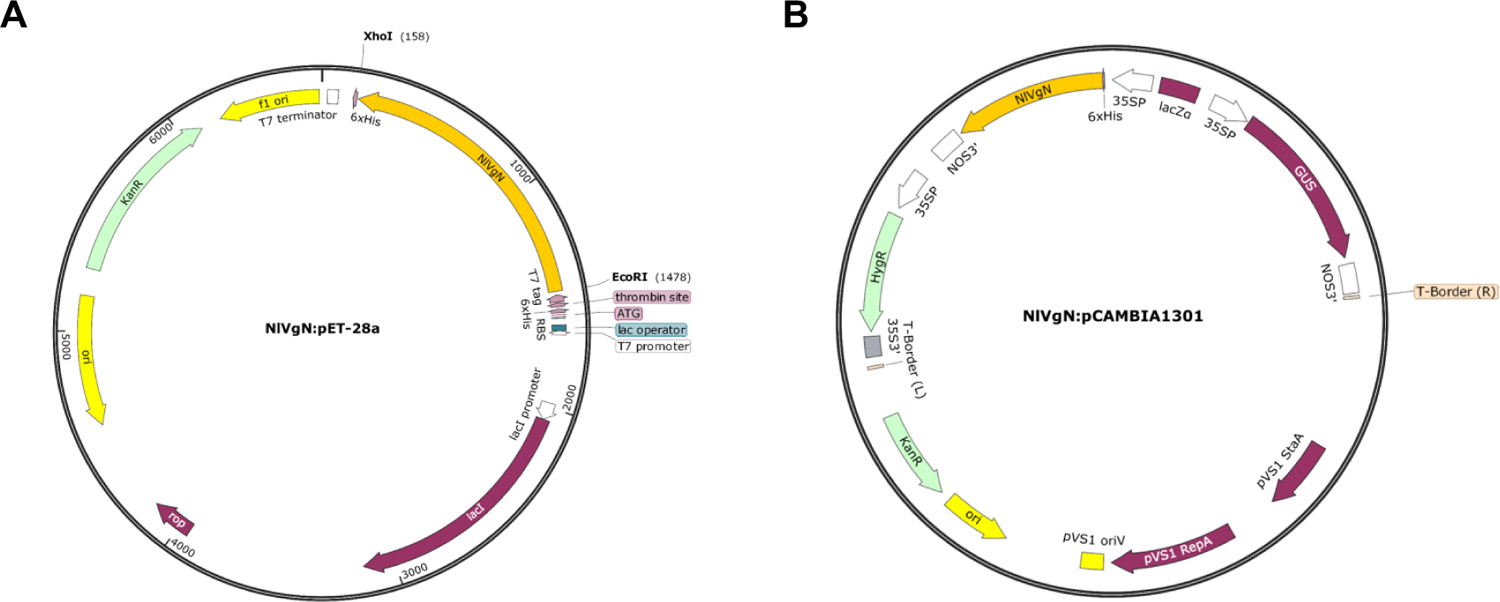
Transformation vectors used for prokaryotic expression of NlVgN or generation of lines expressing *NlVgN*.

